# A parametric texture model based on deep convolutional features closely matches texture appearance for humans

**DOI:** 10.1101/165761

**Authors:** Thomas S. A. Wallis, Christina M. Funke, Alexander S. Ecker, Leon A. Gatys, Felix A. Wichmann, Matthias Bethge

**Affiliations:** Werner Reichardt Center for Integrative Neuroscience, Eberhard Karls Universität Tübingen & the Bernstein Center for Computational Neuroscience, Tübingen; Werner Reichardt Center for Integrative Neuroscience, Eberhard Karls Universität Tübingen, Bernstein Center for for Computational Neuroscience, Tübingen & Department of Neuroscience, Baylor College of Medicine, Houston, TX, USA; Werner Reichardt Center for the Integrative Neuroscience, Eberhard Karls Universität Tübingen & the Bernstein Center for Computational Neuroscience, Tübingen; Neural Information Processing Group, Faculty of Science, Eberhard Karls Universität Tübingen, Bernstein Center for Computational Neuroscience, Tübingen & the Max Planck Institute for Intelligent Systems, Empirical Inference Department, Tübingen; Werner Reichardt Center for Integrative Neuroscience, Eberhard Karls Universität Tübingen, Bernstein Center for Computational Neuroscience, Tübingen, Institute for Theoretical Physics, Eberhard Karls Universität Tübingen & Max Planck Institute for Biological Cybernetics, Tübingen

**Keywords:** spatial vision, natural scenes, texture perception, peripheral vision

## Abstract

Our visual environment is full of texture—“stuff” like cloth, bark or gravel as distinct from “things” like dresses, trees or paths—and humans are adept at perceiving subtle variations in material properties. To investigate image features important for texture perception, we psychophysically compare a recent parameteric model of texture appearance (CNN model) that uses the features encoded by a deep convolutional neural network (VGG-19) with two other models: the venerable Portilla and Simoncelli model (PS) and an extension of the CNN model in which the power spectrum is additionally matched. Observers discriminated model-generated textures from original natural textures in a spatial three-alternative oddity paradigm under two viewing conditions: when test patches were briefly presented to the near-periphery (“parafoveal”) and when observers were able to make eye movements to all three patches (“inspection”). Under parafoveal viewing, observers were unable to discriminate 10 of 12 original images from CNN model images, and remarkably, the simpler PS model performed slightly better than the CNN model (11 textures). Under foveal inspection, matching CNN features captured appearance substantially better than the PS model (9 compared to 4 textures), and including the power spectrum improved appearance matching for two of the three remaining textures. None of the models we test here could produce indiscriminable images for one of the 12 textures under the inspection condition. While deep CNN (VGG-19) features can often be used to synthesise textures that humans cannot discriminate from natural textures, there is currently no uniformly best model for all textures and viewing conditions.

## Introduction

Textures are characterised by the repetition of smaller elements, sometimes with variation, to make up a pattern. Significant portions of the visual environment can be thought of as textures (“stuff” as distinct from “things”; Adelson & Bergen, 1991): your neighbour’s pink floral wallpaper, the internal structure of dark German bread, the weave of a wicker basket, the gnarled bark of an old tree trunk, a bowl full of prawns ready for the barbie. Texture is an important material property whose perception is of adaptive value (Adelson, 2001; Fleming, 2014). For example, we can readily discriminate wet from dry stones (e.g. Ho, Landy, & Maloney, 2008), separating the underlying spatial texture from potentially temporary characteristics like glossiness. Where surfaces of different textures form occlusion boundaries, texture can provide a powerful segmentation cue; conversely, occlusion borders of similarly-textured surfaces can camoflage the occlusion (hiding a tiger among the leaves). Given the importance and ubiquity of visual textures, it is little wonder that they have received much scientific attention, not only from within vision science but also in computer vision, graphics and art (see Dakin, 2014; Landy, 2013; Pappas, 2013; Rosenholtz, 2014, for comprehensive recent reviews of this field).

## Studying texture perception with parametric texture models

Seminal early work on visual texture perception includes that by Gibson (Beck & Gibson, 1955; Gibson, 1950) and by Julesz (Julesz, 1962, 1981; Julesz, Gilbert, & Victor, 1978). Julesz’ thinking remains an important influence on approaches to texture perception, in particular the idea that there exists some set of statistics (parameters in a parametric model) that are both necessary and sufficient for matching the appearance of textures (see also Portilla & Simoncelli, 2000). For computer vision applications, where a goal might be to match the appearance of some region of texture to facilitate image compression, the most effective approaches can be nonparametric—for example, by *quilting* repetitions of a base level crop over the area of the texture (e.g. Efros & Freeman, 2001). However, non-parametric approaches have little to teach us about the human visual system because they make no explicit hypotheses about what features are represented. In this paper we will therefore focus on parametric texture models.

Parameteric models that aim to match the appearance of natural textures are typically assessed by examining artificial textures synthesised by the model (Heeger & Bergen, 1995; Portilla & Simoncelli, 2000; Safranek & Johnston, 1989; Safranek, Johnston, & Rosenholtz, 1990; Zhu, Wu, & Mumford, 1998). The statistics of a model are first computed on a target image, then a new image is synthesised to approximately match the statistics of the target image (often via gradient descent). This approach carries forward Julesz’ “necessary and sufficient statistics” idea by assuming that texture appearance can be captured by the coefficients of some specified set of image statistics. Note that this focus on naturalistic appearance is distinct from a complementary approach which starts from local analysis of luminance distributions to posit an “alphabet” of independent microtexture dimensions (Victor, Thengone, & Conte, 2013), but does not seek to match the appearance of natural textures.

A number of parametric texture models operate by assuming a plausible image representation for the early primate visual system, decomposing the target image into some number of frequency and orientation bands (Cano & Minh, 1988; Heeger & Bergen, 1995; Malik & Perona, 1990; Porat & Zeevi, 1989; Portilla & Simoncelli, 2000; Simoncelli & Portilla, 1998; Zhu et al., 1998). The spatially-averaged responses in some combination of these bands form the parameters of the model, whose values are then matched by the synthesis procedure. The parametric texture model of Portilla and Simoncelli (Portilla & Simoncelli, 2000; Simoncelli & Portilla, 1998) extended this approach by additionally matching the correlations between channels and other statistics, producing more realistic appearance matches to textures. This model has since had broad impact on the field of human perception and neuroscience: the texture statistic representation may provide a fruitful way to understand the processing in mid-ventral visual areas (Freeman & Simoncelli, 2011; Freeman, Ziemba, Heeger, Simoncelli, & Movshon, 2013; Freeman, Ziemba, Simoncelli, & Movshon, 2013; Movshon & Simoncelli, 2014; Okazawa, Tajima, & Komatsu, 2015; Ziemba, Freeman, Movshon, & Simoncelli, 2016), and it has been argued to provide a good approximation of the type of information encoded in the periphery, and thus a model for tasks such as crowding and visual search (Balas, Nakano, & Rosenholtz, 2009; Freeman & Simoncelli, 2011; Keshvari & Rosenholtz, 2016; Rosenholtz, 2011; Rosenholtz, Huang, & Ehinger, 2012; Rosenholtz, Huang, Raj, Balas, & Ilie, 2012).

How, though, does it perform as a model of texture appearance in humans? Balas and colleagues (Balas, 2006, 2008, 2012; Balas & Conlin, 2015) have reported a number of psychophysical investigations using the Portilla and Simoncelli (hereafter, *PS*) texture model that are relevant to this question. Balas (2006) quantified the relative importance of subsets of the PS statistics compared to the full set for matching the appearance of different classes of texture (periodic, structured or asymmetric). He used a task in which human observers chose the “oddball” image from a set of three (a three-alternative oddity task) that were presented briefly to the near-periphery. Two of the images were drawn from original textures whereas the oddball was drawn from a model synthesis matched to the original texture (or vice versa; the oddball could be either original or synthetic). Importantly, all three images were physically different from each other (consisting of subcrops of larger images). The oddity judgment therefore concerns the subjective dissimilarity of the images—which image is “produced by a different process”—rather than exact appearance matching. In this study, the importance of different parameter subsets depended on the class of texture, and including the full set of statistics brought average discrimination performance quite close to chance (around 40% correct on average), showing that the PS statistics do a reasonably good job in capturing texture appearance under brief peripheral viewing conditions.

Balas (2012) used a four-alternative oddity task to investigate the discriminability of real and synthetic textures. Observers were allowed to view each stimulus array for unlimited time and to foveate the images. Under these viewing conditions, observers could easily discriminate original natural textures and PS-synthesised images from each other, whether the oddball was real or synthetic (average performance 85–90%). However, when the original images were sourced from abstract artworks rather than photographs of fruits and vegetables, performance for discriminating real from PS-synthesised images was worse and depended on whether the oddball was real or synthesised (with performance around 55% for the former and 65% for the latter). Together with the results of Balas (2006), these results suggest that the PS model better captures texture appearance in the periphery than in the fovea, and that the perceptual fidelity of the matching depends on the image or texture type.

Finally, Balas and Conlin (2015) assessed whether the influence of illumination change on human texture perception could be captured by PS synthesis. Observers performed a match-to-sample task, in which they decided which of two match images depicted the same texture as a previously-presented sample. Performance was quite high (above 90%) when the illumination between the sample and correct match image was constant (in this case, the match image was physically identical to the sample), whether the images were real or synthesised. When the correct match image was presented with different illumination to the sample, performance declined to around 70% correct for synthetic images but remained high for real images. That is, observers could easily ignore illumination changes when matching real textures, but their judgments were impaired by illumination change when discriminating synthesised images. Note that the foil images (the non-target match image) were selected to be “approximately visually-matched” by the experimenters; it is likely that the results (but perhaps not conclusions) will depend on this choice. Similar results were obtained after equalising the luminance and power spectra of the images, and when match and sample images were physically different (cropped from different areas of the same texture). These results show that the PS feature space does not perfectly preserve the necessary statistics to match texture appearance across changes in illumination.

Together, the experiments show that while aspects of human texture perception are not captured by or fall outside the scope of the PS feature space, it does succeed in capturing key aspects of texture appearance for many classes of natural texture. The PS feature space is based on the idea—amply supported by psychophysical and neurophysiological evidence—that the human visual system decomposes an image into a number of spatial and orientation subbands. To what extent will a more complex feature space improve on the PS model?

## A new parametric texture model based on deep features

Gatys et al. (2015) have recently introduced a new parametric texture model that produces subjectively high-quality matches to texture appearance, and whose features can be used to separate the “style” of an image from its content (Gatys, Ecker, & Bethge, 2016). This texture synthesis procedure (see “CNN texture model”, below) is based on the pre-trained features of a deep convolutional neural network (the VGG-19; Simonyan & Zisserman, 2015, Figure 1) that achieves near state-of-the-art performance on the Imagenet Large Scale Visual Recognition Challenge (ILSVRC; Russakovsky et al., 2015): basically, returning labels for the likely objects present in an image. Due to their success on benchmarks like the ILSVRC, CNNs have become the dominant approach to many visual inference problems in the field of computer vision, with some networks showing impressive transfer learning performance (doing well on new tasks with only minimal changes to the network, e.g. Donahue, Jia, & Vinyals, 2013).

**Figure 1.**
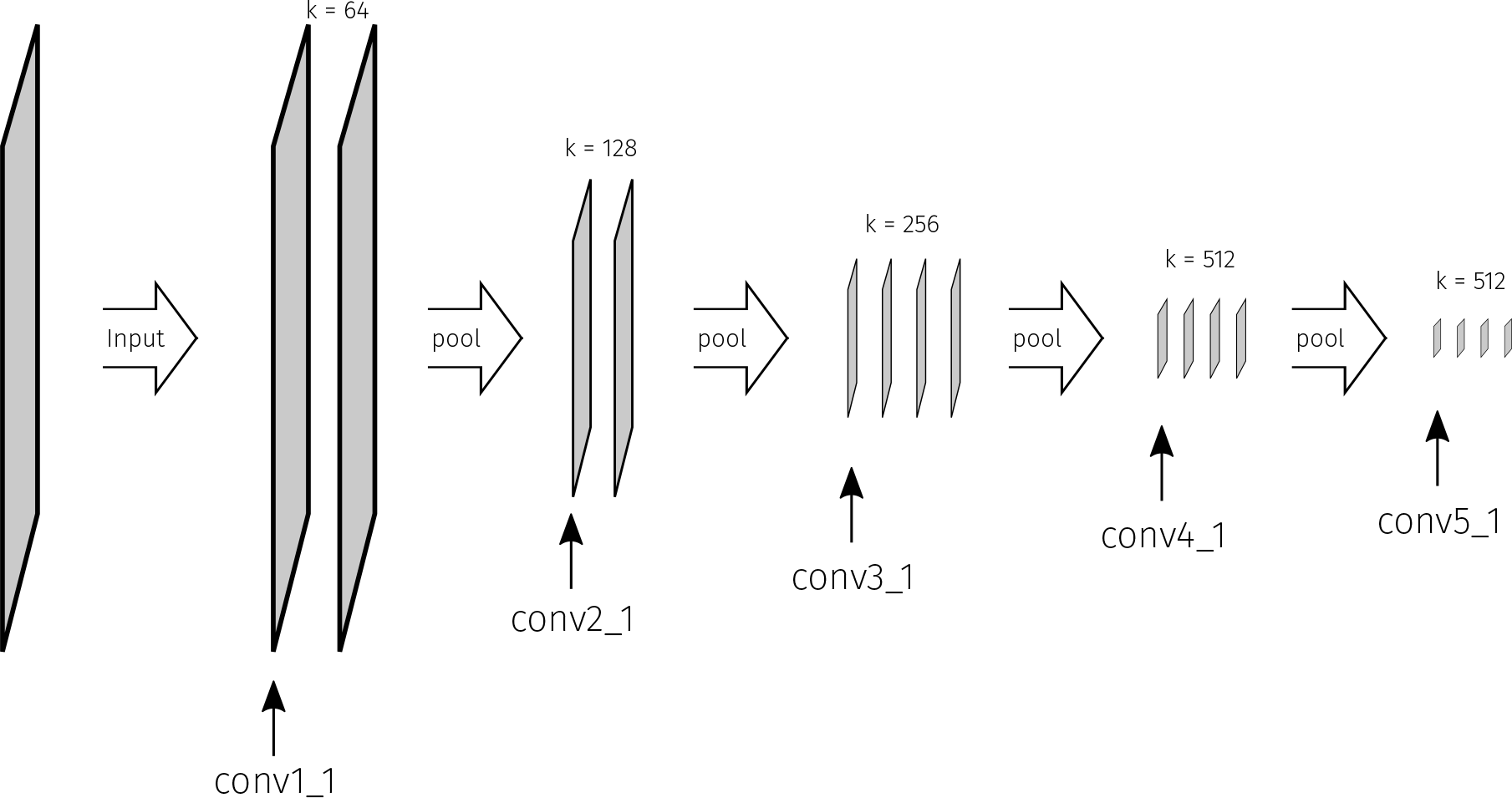
The architecture of the VGG-19 convolutional neural network (Simonyan & Zisserman, 2015), whose pretrained features are used by the Gatys, Ecker, and Bethge (2015) texture model. The network consists of stacks of convolutional stages followed by max-pooling. In higher network layers, the feature map sizes decrease (depicted as the increasingly small panels), the corresponding “receptive field” sizes of the units increase, and the number of feature maps (*k*) increase. In this paper we synthesise textures using the first convolutional layer from each stack after the max pooling.

Briefly, a single-layer convolutional neural network (CNN) learns (via supervised training) the weights of filters that are convolved with input images, creating a spatial feature map of activations, similar to a traditional bank of Gabor filters familiar to vision scientists. Using convolutional filters allows the detection of spatial patterns at any position in the image (translation equivariance), and also facilitates learning through weight sharing—the intuition here is that features useful to know about at one spatial location are likely to be useful for all spatial locations. All convolutional layer activations are then passed through a pointwise nonlinearity, typically a rectified linear (“relu”) function *f* (*x*) = max(0, *x*). These feature maps can then be pooled (in VGG by taking the maximum of activations in a small area), creating local spatial invariance, and combined with downsampling to reduce the spatial dimensions of the feature maps (see Figure 1). Stacking such operations repeatedly (passing the outputs of one convolutional or max-pool layer as the input to another, creating a “deep” CNN with at least one hidden layer) has several effects. The spatial area of the input image to which features respond are larger for higher layers (analogous to the increase in receptive field size from V1 to IT cortex), and the features to which higher convolutional layers respond becoming increasingly non-linear functions of the input pixels (analogous to the feature selectivity from V1 to IT cortex). It is this accumulating nonlinear behaviour that allows complex properties such as object identity (and many other properties; Hong, Yamins, Majaj, & DiCarlo, 2016) to be linearly decoded from the higher network layers. For more comprehensive recent reviews, see Kietzmann, McClure, and Kriegeskorte (2017); LeCun, Bengio, and Hinton (2015) and Yamins and DiCarlo (2016).

CNNs are interesting for the study of human vision first and foremost because they perform interesting tasks. Until recently, there was only one known class of system (“biological brains”) that could detect and recognise objects in photographic images with high accuracy; now there are two. The second reason that human vision researchers might be curious about CNNs is that there is growing evidence that the *way* in which CNNs perform these tasks has intriguing similarities to some biological visual systems. For example, there is now quantitative evidence that performance-optimized CNN features predict ventral stream brain signals in monkeys and humans using the stimulus input better than existing models built explicitly for that purpose (Cadieu et al., 2014; Cichy, Khosla, Pantazis, & Oliva, 2016; Cichy, Khosla, Pantazis, Torralba, & Oliva, 2016; Guclu & van Gerven, 2015; Hong et al., 2016; Khaligh-Razavi & Kriegeskorte, 2014; Yamins, Hong, Cadieu, & DiCarlo, 2013; Yamins et al., 2014). CNN models also show similarities to human psychophysical object recognition performance under brief presentation conditions (Hong et al., 2016; Yamins et al., 2014). A recent paper reported that CNNs trained on ImageNet (natural photos) can still partially recognize objects from silhouette information only, and show other human-similar shape biases (Kubilius, Bracci, & Op de Beeck, 2016). There are of course important ways that current CNNs are *un*like primate visual systems. For example, a subtle modification of an image that is nearly imperceptible to a human can cause a deep network to misclassify an object with high confidence (Szegedy et al., 2013, see Yamins and DiCarlo (2016) for additional discussion). Furthermore, human object recognition remains remarkably robust in images degraded by white noise, whereas the original VGG network is strongly impaired (Geirhos et al., 2017). Bearing these caveats in mind, an exciting possibility is that the study of CNNs may help to elucidate some fundamental mechanisms of human perception.

In this paper we pursue a less lofty goal: to measure how well humans can discriminate textures synthesised by the Gatys et al. (2015) model from natural textures. How well do CNN texture features match the appearance of the original textures? To address this question we compare the model of Gatys et al. (2015) to the PS model (Portilla & Simoncelli, 2000) and to a recent modification of the Gatys model (Liu, Goussau, & Xia, 2016). Experimentally, we closely follow the approach of Balas (2006), described above^
1
^. Using images that are all physically different measures the extent to which model syntheses are *categorically* or *structurally* lossless (in that they could both be considered samples from original images; Pappas, 2013), as opposed to being *perceptually* lossless (unable to be told apart) compared either to each other (Freeman & Simoncelli, 2011) or the original source images (Wallis, Bethge, & Wichmann, 2016). Perceptual losslessness could be important for understanding visual encoding in general but categorical losslessness is arguably more useful for understanding the perceptual representation of texture.

In addition to assessing the discriminability of brief, peripherally-presented textures (as in Balas, 2006), we are also interested in how this changes when longer foveal comparison is possible (as in Balas, 2012). We therefore include two presentation conditions: a “parafoveal” condition and an “inspection” condition^
2
^. Note that depending on the spatial scale of the most informative differences, sensitivity to some aspects of texture can be *better* in the parafovea than in the fovea under some conditions (Gurnsey, Pearson, & Day, 1996; Kehrer, 1987, 1989). Therefore, differences in psychophysical performance between these conditions are informative about the extent to which the texture models under consideration capture, or fail to capture, features that are important for both foveal and near-peripheral texture perception.

## General methods

All stimuli, data and code to reproduce the figures and statistics reported in this paper are provided online (Raw data and code at http://doi.org/10.5281/zenodo.438029, stimuli at http://doi.org/10.5281/zenodo.438031). This document was prepared using the knitr package (Xie, 2013, 2015) in the R statistical environment (Arnold, 2016; Auguie, 2016; {R Core Development Team}, 2016; Wickham, 2009, 2011; Wickham & Francois, 2016) to improve its reproducibility.

### Apparatus

Stimuli were displayed on a VIEWPixx 3D LCD (VPIXX Technologies; spatial resolution 1920 × 1080 pixels, temporal resolution 120 Hz, operating with the scanning backlight turned off in high-bitdepth greyscale mode). Outside the stimulus image the monitor was set to mean grey. Observers viewed the display from 60 cm (maintained via a chinrest) in a darkened chamber. At this distance, pixels subtended approximately 0.024 degrees on average (41 pixels per degree of visual angle). The monitor was linearised (maximum luminance 260 cd/m^
2
^) using a Konica-Minolta LS-100 photometer. Stimulus presentation and data collection was controlled via a desktop computer (Intel Core i5-4460 CPU, AMD Radeon R9 380 GPU) running Ubuntu Linux (16.04 LTS), using the Psychtoolbox Library (Brainard, 1997; Kleiner, Brainard, & Pelli, 2007; Pelli, 1997, version 3.0.12) and our internal iShow library (http://dx.doi.org/10.5281/zenodo.34217) under MATLAB (The Mathworks, Inc., R2015b).

### Source images

Twelve unique texture images^
3
^ (see Figure 2) were selected to provide avariety of texture-like structure (including some with obvious periodicity and others that were asymmetric) but were also chosen to exhibit some non-texture naturalistic structure (such as the size gradient visible in the flowerbed image). Images were converted to greyscale using scikit-image’s io.imread function (van der Walt et al., 2014), then cropped to the largest possible square from the centre of the image. The original images all had at least one dimension of 1024 pixels. We then downsampled all images to 256 × 256 pixels using the cubic interpolation of skimage.transform.resize. To preserve the naturalistic appearance of the images we did not standardise the mean or variance of intensities. Since all texture models considered here also match the low-level image statistics this will not impact our results. For each image model (conv1–conv5 and PS for Experiment 1, conv5, PS and powerspec in Experiment 2; see below) we generated ten unique synthesised images of size 256 from each original image, resulting in a final stimulus set of 732 images for Experiment 1 and 372 images for Experiment 2. All images were stored as 16-bit .png files.

**Figure 2.**
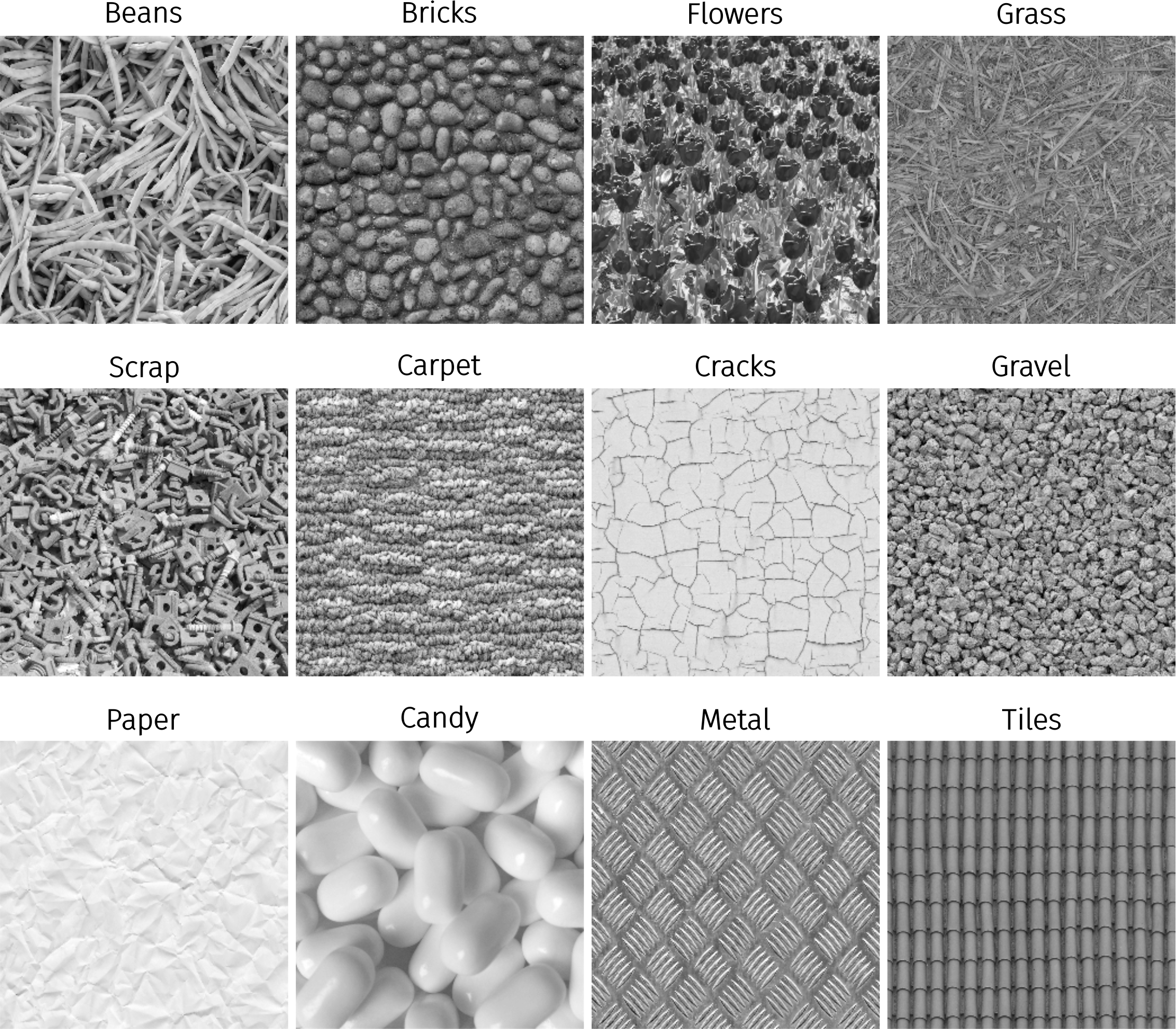
The twelve original texture images used in the experiments. Arranged to correspond to Figure 7. These images are copyrighted by www.textures.com (used with permission).

### CNN texture model

The CNN texture model (Gatys et al., 2015) uses the pre-trained features of the VGG-19 network (Simonyan & Zisserman, 2015), which shows near state-of-the-art performance on the object recognition ImageNet challenge (Russakovsky et al., 2015). While there are now CNN models that outperform the VGG network on object recognition, the VGG network remains appealing because of its relatively simple architecture (Figure 1), and because it produces more introspectively appealing textures and style transfer than those networks that currently performing better on ImageNet. It consists of two operations, stacked many times: convolutions with *k* 3 × 3 filters (where *k* is the number of input feature maps) followed by a 2 × 2 max-pooling in non-overlapping regions. The model uses five pooling and 16 convolutional layers (plus three fully-connected layers which we do not use here). The layers are typically labelled with the stack (e.g. “conv1” or “pool1”) with an underscore denoting the sub-layer. For example, “conv1_1” refers to the first convolutional layer of the network, whereas “conv3_2” would be the second convolutional layer of the third stack (Figure 1). We use a subset of these feature maps for texture synthesis (see below). The code was implemented in Theano using the Lasagne framework, and may be downloaded from https://github.com/leongatys/DeepTextures. The weights of the VGG-19 network are scaled such that the mean activation of each filter over images and positions is equal to 1.

The first step of the texture synthesis algorithm is to pass the original image through the network, generating responses in all network layers. For the feature responses of a subset of layers (described below) the Gram matrices are computed (the Gram matrix is the dot product of the vectorised feature maps; each entry in the resulting matrix is the correlation between two features in response to a particular input image). The basic idea of the texture synthesis is to create an image with the same Gram matrix representation via gradient descent (the same synthesis principle as in Portilla and Simoncelli (2000) using different features). We start with a white noise image and minimize the mean-squared distance between the entries of the Gram matrices of the original image and the Gram matrix of the image being generated. For the optimization we use the L-BFGS method from the the SciPy package (Jones, Oliphant, & Peterson, 2001) using 1000 iterations, which was sufficient to bring the loss to an acceptable (but usually nonzero) value. Note that this procedure (using a unique random initialisation and converging on nonzero loss) can therefore generate an effectively infinite number of physically unique synthesised images. We discuss the gradient descent further in the Appendix. After gradient descent, the intensity histogram of the resulting image was matched to the intensity histogram of the original image (ensuring that the images have the same global luminance, contrast, skew and kurtosis).

The network was trained on RGB images and expects three-channel input. We duplicated the greyscale original images into three channels, and to ensure that the outputs of the synthesis remained greyscale we averaged the gradients of each colour channel during optimization. The layers conv1_1, conv2_1, conv3_1, conv4_1 and conv5_1 were used for texture synthesis by taking the activations after rectification. For simplicity, we label the texture models used below with the name of the highest convolutional stack used. We match all the Gram matrices cumulatively up to the named layer (e.g. the model we label “conv3” below matches Gram matrices for layers conv1_1, conv2_1 and conv3_1). For each layer *l* with *n_l_
* feature maps, 
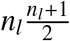
 parameters are matched (division by two is because the Gram matrices are symmetrical). The approximate number of parameters in each CNN texture model are shown in Figure 5. Outputs were saved as 16-bit .png images. Example syntheses can be seen in Figure 3.

**Figure 3.**
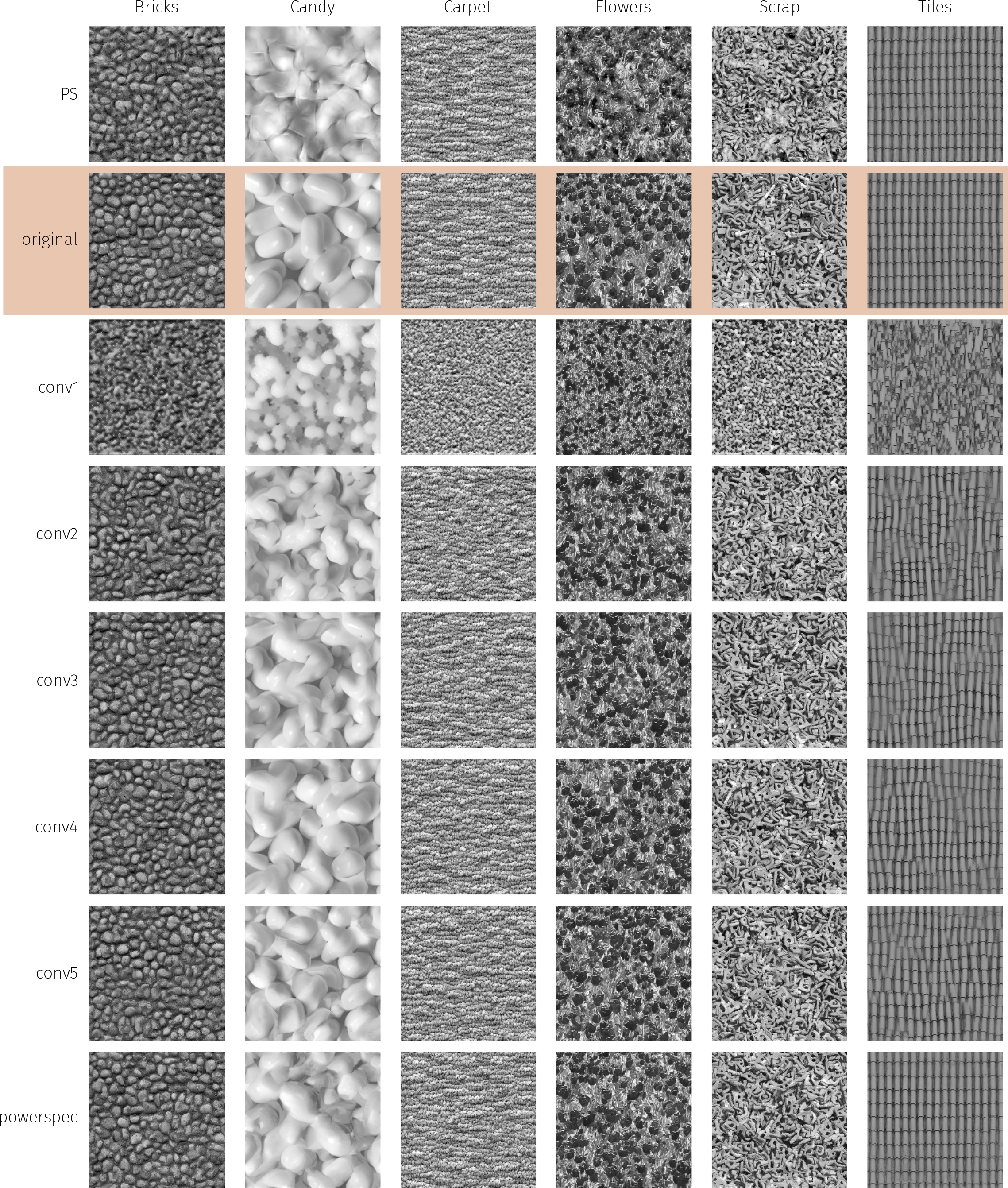
Example experimental stimuli used in Experiment 1 (PS, conv1–conv5) and Experiment 2 (PS, conv5 and powerspec).

**Figure 5.**
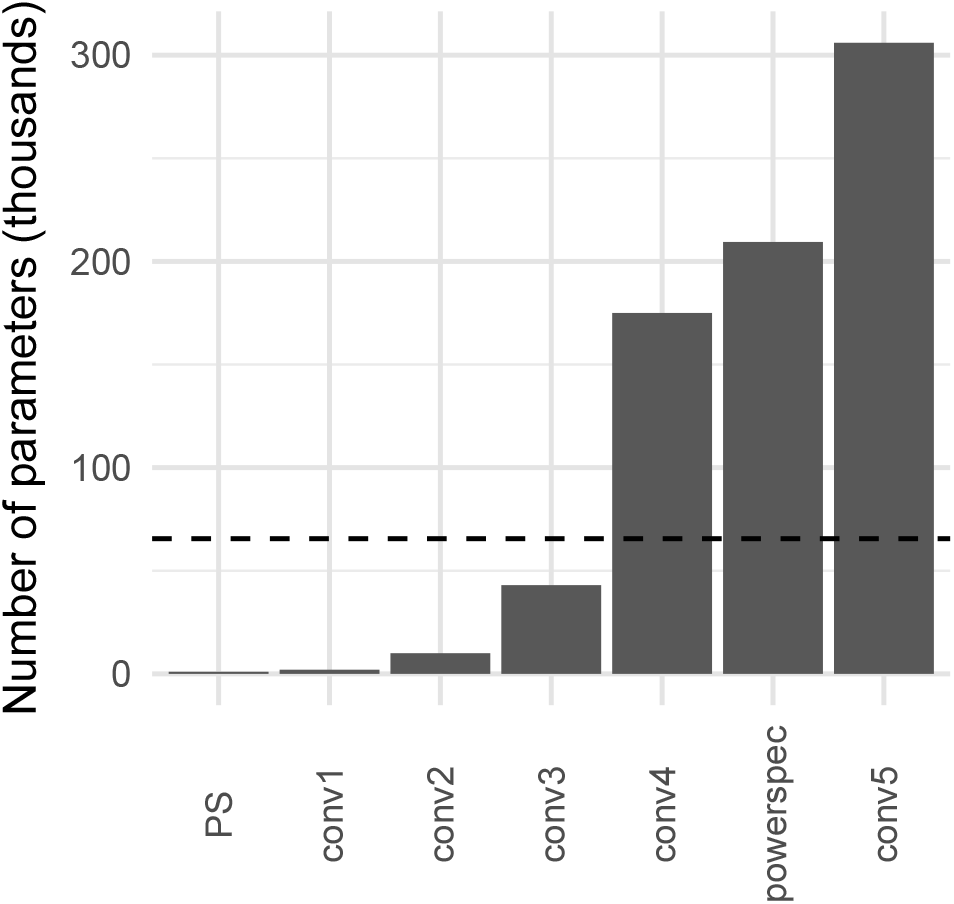
Approximate number of parameters matched by each texture model assessed in the present paper. The dashed line shows the number of pixels in a 256-pixel-square image. Models above the line are over-complete.

### CNN plus power spectrum model

To capture long-range correlations (such as contours that extend over large sections of the image) the model can be extended by additionally matching the power spectrum of the original image when performing the gradient descent to find texture syntheses (Liu et al., 2016). The new loss function is *L* = *L*
_CNN_ + *βL*
_spe_ and the new gradient is Δ = Δ_CNN_ + *β*Δ_spe_, where *L*
_CNN_ is the loss function and Δ_CNN_ is the gradient from the pure CNN texture model, *L*
_spe_ and Δ_spe_ are related to the distance between the current image and the target Fourier spectrum, and *β* = 10^
5
^. That is, the additional constraints are simply added into the loss function and gradient (see Liu et al., 2016, for further details).

To synthesise these stimuli we used code provided by Liu et al. (2016). There are a number of differences between the implementation of the power spectrum model and the base CNN model described above. First, the code is written using Matconvnet instead of Lasagne. The network and the images are normalised to [0, 1] and the stopping criterion of the optimisation process is different. In the powerspectrum model we used up to 2000 iterations (as distinct to 1000 iterations for the base model). The powerspectrum model matches different layers of the VGG compared to our CNN model: Conv1_1, Pooling1, Pooling2, Pooling3 and Pooling4. The power spectrum constraint adds 32,768 parameters (half the size of the image because phase is discarded), yielding a total of 209,408 parameters (Figure 5). While we have not run extensive experiments, we argue that the most consequential change between the models for the results we report is the inclusion of the powerspectrum matching constraint rather than other implementation differences.

### PS texture model

Portilla and Simoncelli (PS Portilla & Simoncelli, 2000) texture images were generated using the publically-available MATLAB toolbox (http://www.cns.nyu.edu/lcv/texture/). The texture synth representation we used consisted of four spatial scales and orientations, and a spatial neighbourhood of 11 pixels (these are the most common settings used in the literature where reported (e.g. Balas et al., 2009; Freeman & Simoncelli, 2011)). The gradient descent procedure was based on 50 iterations. The PS model matches approximately 1,000 parameters (Figure 5). Outputs were saved as 16-bit .png images.

### Procedure and design

On each trial observers were presented with three physically different image patches. Two were from the original image and one from a model synthesis image matched to that original image (or vice versa—two patches could come from the same model synthesis and one patch from the original image). That is, the odd-ball image could be either original or synthesised with equal probability, so a “pick the natural-looking image” strategy would not succeed. The three image patches (size 128-pixels-square) were cropped from a larger image (size 256). To obtain two non-overlapping crops from the same physical image (for the nontarget intervals) one could simply use the image quadrants. To increase the physical variation in the images across trials we instead chose two adjacent crops drawn from non-overlapping but otherwise jittered image sections. On half of the trials the crops were from adjacent “rows” with the vertical dimension randomly sampled, whereas on the other half the crops were from adjacent columns with horizontal dimension randomised. This strategy eliminated the possibility that observers could match specific features of the images within a trial (as in Balas, 2006).

The oddball image could appear at any one of three locations with equal probability (see Figure 4). The observers’ task was to report which of three simultaneously-presented images was different to the other two, in that it was “generated by a different process” (rather than being physically the same). Observers fixated a spot (best for steady fixation from Thaler, Schütz, Goodale, and Gegenfurtner (2013)) in the centre of the screen, and the images were arranged around the fixation in a downward-pointing equilateral triangle configuration. The images were windowed by a circular cosine, ramping the contrast to zero in the space of 6 pixels. The distance between the fixation point and the nearest edge of the image was 3 degrees of visual angle (dva), and the image patches subtended 3.1 dva.

**Figure 4.**
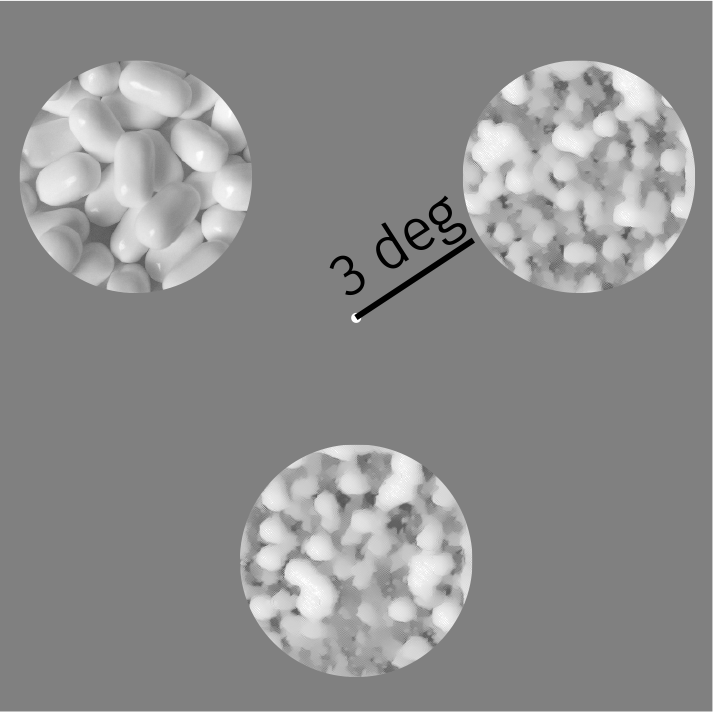
Example experimental display (not to scale). Distance bar not shown in actual experiment.

The stimulus display was presented for either 200 ms, with observers instructed to maintain fixation (the *parafoveal* condition) or for 2000 ms with observers allowed to make eye movements freely (the *inspection* condition). Observers then had 1200 ms to respond (responses could also be made while the stimulus remained on the screen). The inter-trial interval was 400 ms. To reduce the possibility that observers could learn specific strategies for different images based on familiarity, no trial-to-trial feedback was provided. Instead, a break screen was presented every 72 trials telling the observer their mean performance on the previous trials.

Within a block of trials observers saw five repetitions of the 72 combinations of image model (six levels) and source image (12 levels), for a total of 360 trials per block. Trials were pseudo-randomly interleaved throughout a block, with the constraint that trials using the same source image were required to be separated by at least two intervening trials. Presentation condition was blocked to allow observers to anticipate the trial timing and adjust their strategy accordingly.

At the beginning of the experiment, naïve observers performed 30 trials with a 2 second presentation time to allow them to become familiar with the task. All observers then performed a practice session of 30 trials at the relevant presentation time for the upcoming block.

### Data analysis

We analysed the data using a logistic Generalised Linear Mixed Model (GLMM), estimated using Bayesian inference. Experimentally-manipulated fixed effects of presentation condition and image model were estimated along with random effects for observer and image. The model parameters were given conservative, weakly-informative prior distributions such that we assumed no effects of our experimental manipulations (by using priors for regression parameters centred on zero) but with high uncertainty. This biases the model against finding spuriously large effects. Bayesian model estimation offers two practical advantages here: first, posterior credible intervals over model parameters have an intuitively-appealing meaning (they represent our belief that the “true” parameter lies within some interval with a given probability, conditioned on the priors, model and data). Second, the priors act to sensibly regularise the model estimates to ensure all parameters are identifiable. More details and analysis are provided in the Appendix.

## Experiment 1: Original texture model

This experiment compares textures produced by the CNN texture model to the PS model under two observation conditions. This experiment was conducted on two groups of observers. The first (Experiment 1a) consisted of two of the authors, who were familiar with the stimuli, experienced with the psychophysical task, and optically corrected as appropriate. The authors completed five experiment sessions (each consisting of one parafoveal and one inspection block), for a total of 3600 trials each. The order of presentation conditions was pseudorandomly determined for each author in each experiment session. The dataset consisted of 7200 trials.

The second group (Experiment 1b) consisted of ten^
4
^naïve observers (median age 25 years, min 21, max 36), who completed only one experimental session each (i.e. one block of each presentation time)^
5
^. They were paid 10 EUR for the one hour session. Half the observers saw the parafoveal condition first, whereas the other half performed the inspection condition first. All protocols conformed to Standard 8 of the American Psychological Associations Ethical Principles of Psychologists and Code of Conduct (2010) and to the Declaration of Helsinki (with the exception of article 35 concerning pre-registration in a public database). The final dataset consisted of 7200 trials.

### Results

Performance as a function of image model and presentation time, averaging over images, is shown in Figure 6. More complex CNN models (matching more parameters) tend to produce poorer psychophysical performance (i.e. better matches to natural appearance), and the performance in the parafoveal condition is poorer than the inspection condition. The PS model produces better psychophysical performance (i.e. is not as good at matching appearance) than the higher-layer CNN models under the inspection condition but not under the parafoveal condition. The average pattern of results for the ten naïve observers is qualitatively similar to the data shown by the two authors, with the exception that performance is slightly lower. The figure additionally demonstrates what might be believed about the “population of texture images” from our results. Estimates and credible intervals from the mixed-effects model are shown as lines and shaded areas in Figure 6 (further details and quantification are provided in the appendix).

**Figure 6.**
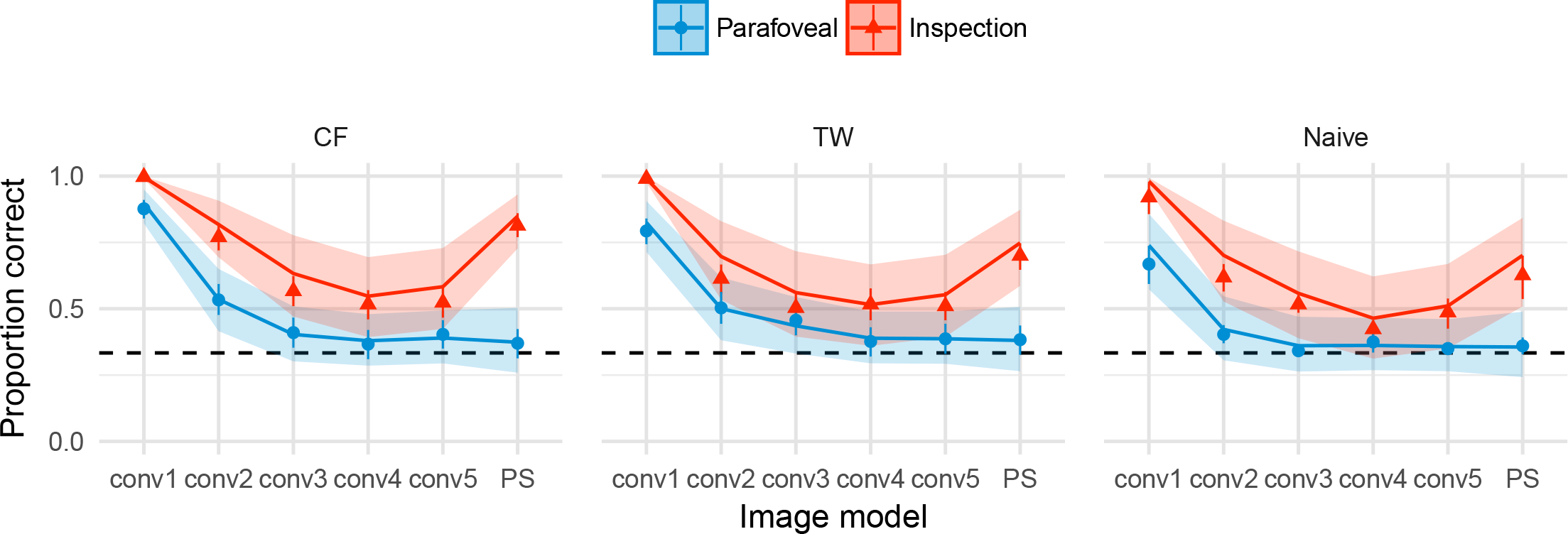
Performance as a function of image model in Experiment 1, averaging over images. For the authors (CF and TW), points show the mean proportion correct and error bars show 95% bootstrapped confidence intervals. Each data point represents 300 trials. Solid lines show mixed-effects model predictions for this observer (mean of posterior regression line), ignoring random effects of image. For naive observers (right panel, N = 10), points show grand mean and 95% bootstrapped confidence intervals based on individual observer means; lines represent mixed-effects model predictions and uncertainty for the population fixed effects, ignoring random effects of observer and image. The dashed horizontal lines in all panels show chance performance. Shaded regions in all panels show 95% credible intervals for the given model. Note these are independent, and so overestimate the uncertainty for making any pairwise comparison between conditions (see appendix for details).

We observe distinctly different effects of image model and presentation time at the level of individual images (Figure 7). Five images (Beans, Bricks, Flowers, Grass and Scrap) show a similar pattern of results as in the average data. Unlike the first five images, the PS model also succeeds in matching appearance for Carpet, Cracks, Gravel and Paper under the inspection condition. In addition, for these images there is less evidence of a difference between the parafoveal and inspection conditions after the conv1 model. These results suggest these four images are easier for all models to synthesise than the first five images. Conversely, all models fail to match the appearance of Metal and Candy under the inspection condition (psychophysical performance well above chance), whereas the parafoveal condition has a marked effect such that performance drops nearly to chance for the higher convolutional and PS models. Finally, the Tiles image is interesting because here the PS model produces better matches to appearance than the CNN models (the syntheses are more difficult to discriminate).

**Figure 7.**
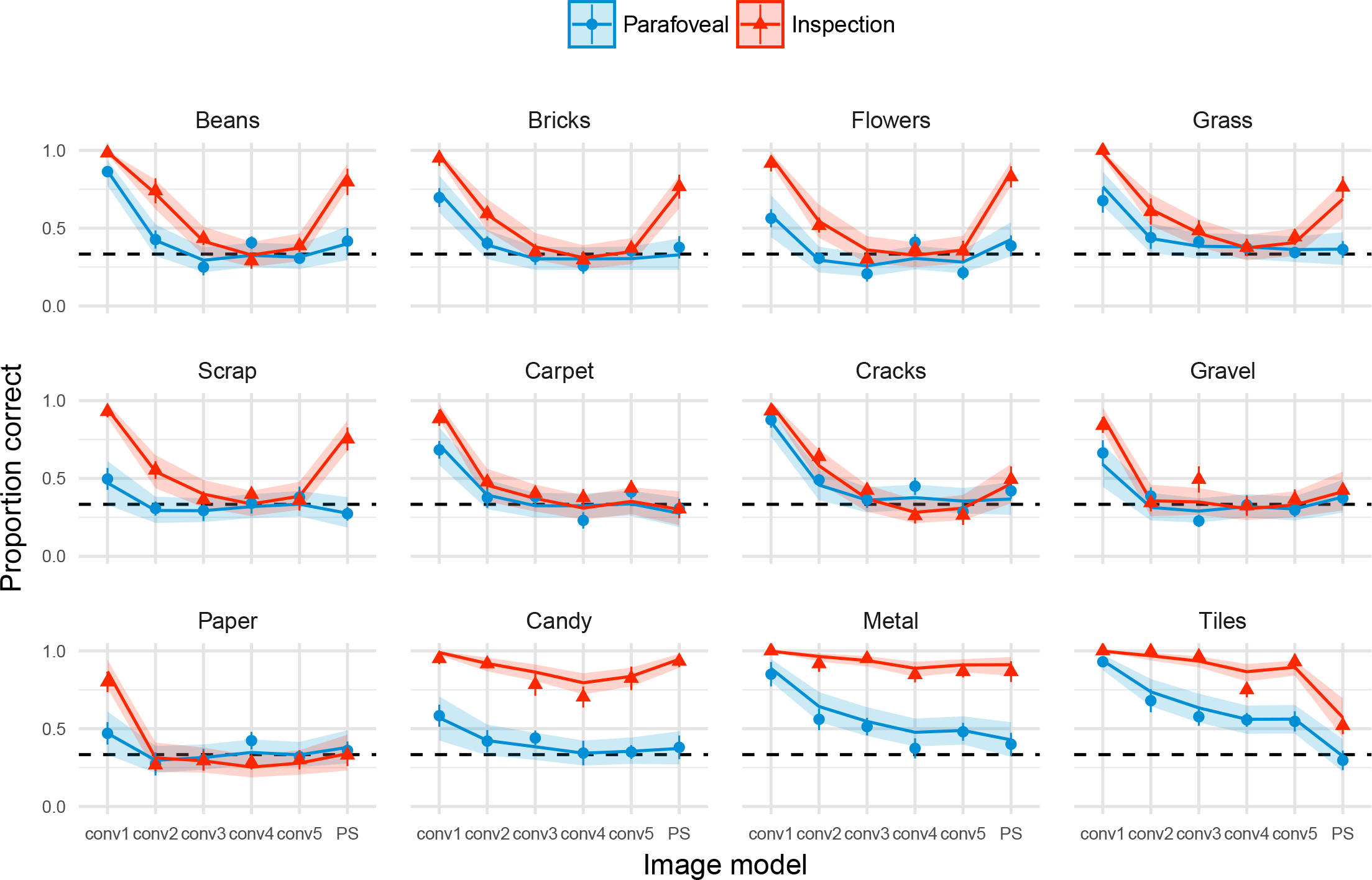
Performance for each image in Experiment 1. Points show the grand mean of all observer means (based on 25 trials for authors and 5 trials for naives). Error bars on points show _1 SEM. Lines show mixed-effects model estimates (posterior mean, including random effects of image but excluding random effects of subject) and shaded regions show 95% credible intervals. That is, the model predicts mean performance to lie in the shaded area with 95% probability, if the image was shown to an average, unknown subject. Images have been arranged according to consistent patterns of results (reading left-to-right). The original images can be seen in Figure 2.

### Experiment 2: Power spectrum constraint

In Experiment 1, the CNN texture model failed to match textures that could be considered “quasi-periodic”, in that they contain global regularities spaced across the whole texture image (for example, the roof tiles or the metal floor textures). Liu et al. (2016) recently showed that such textures can be more closely modelled by adding a power spectrum constraint to the synthesis procedure in CNN texture models. That is, the gradient descent procedure now aims to match both the CNN features and the global Fourier power spectrum of the original image. In an image like the Tiles, the periodic regularity shows up as a strong orientation-and-frequency component in the power spectrum. Matching this improves the perceptual quality of such textures (see Figure 8). In this experiment we seek to quantify this improvement with respect to the unconstrained conv5 model and the PS model for our 12 texture images, using the same procedure as in Experiment 1.

**Figure 8.**
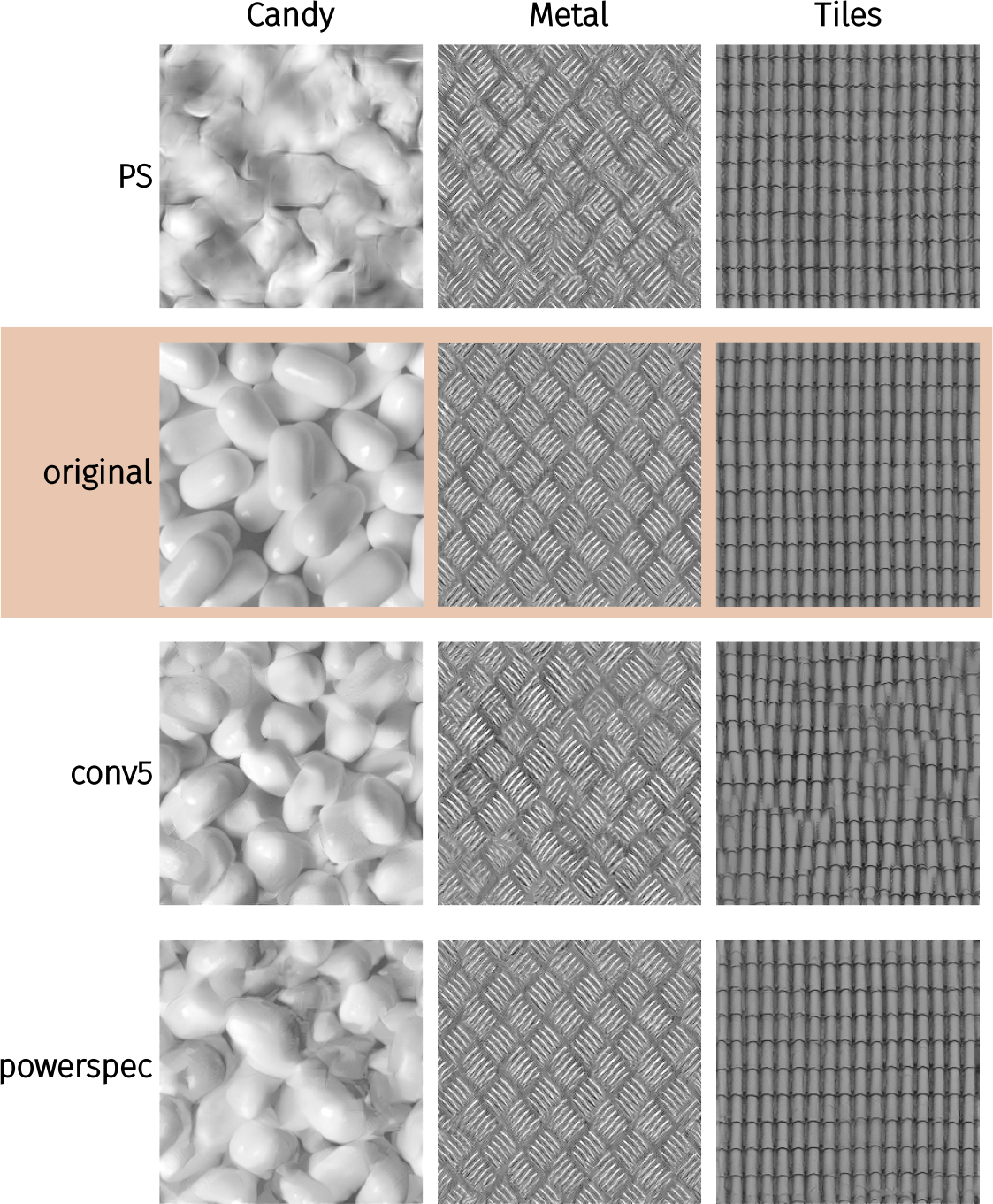
Example experimental stimuli, Experiment 2. Original texture images are copyrighted by www.textures.com (used with permission).

Five observers participated in this experiment, consisting of two authors (CF and TW) and three naïve observers, one of whom had participated in the first experiment. All observers completed two experiment sessions (each consisting of one parafoveal and one inspection block) for a total of 1440 trials, with the exception of S1, who did not return for a second session of testing and so completed only 720 trials.

### Results

For average performance over images (Figure 9) and at the individual image level (Figure 10), the results of Experiment 2 are similar to those of Experiment 1 for the conv5 and PS models. The powerspec model produces similar performance to the conv5 model for most images, with the possible exceptions of Beans, Bricks, Flowers and Grass, in which human performance is slightly higher than for conv5 (i.e. the powerspec model is less effective at matching appearance than conv5). For images with significant long-range regularities (Metal and Tiles) whose appearance failed to be matched by conv5, the powerspec model drastically reduced psychophysical performance. That is, the model syntheses are now approximately matched to the visual appearance of these original images even under foveal inspection (see Appendix). Note however that one observer (author TW) still achieves high accuracy for the powerspec model of Metal, showing that the model fails to capture some important features that at least one observer can see. Finally, all models fail to capture the appearance of the Candy image under inspection.

**Figure 9.**
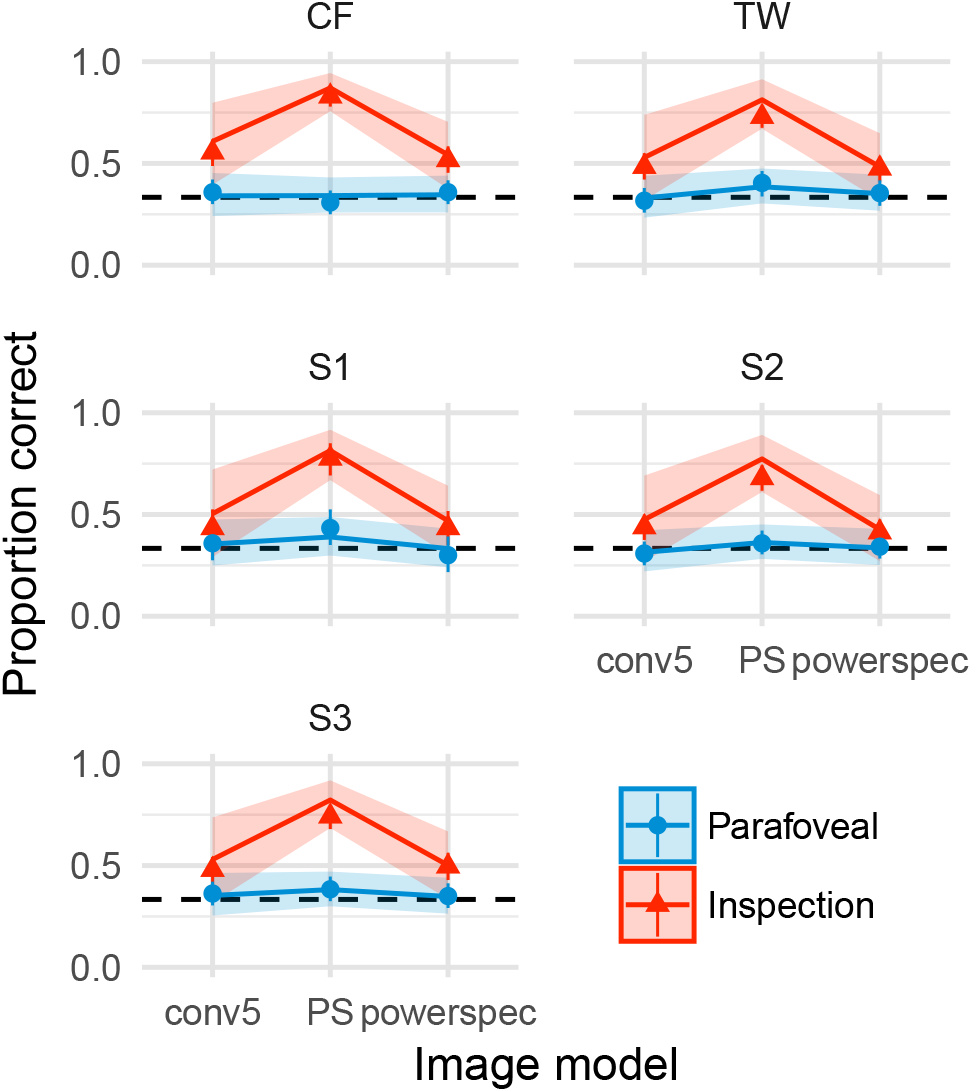
Performance as a function of image model in Experiment 2, averaging over images. Points show mean and 95% confidence intervals on performance (each based on 240 trials for all observers except S2). Lines show mixed-effects model predictions for each observer (mean of posterior regression line) and shaded regions show model 95% credible intervals, ignoring random effects of image.

**Figure 10.**
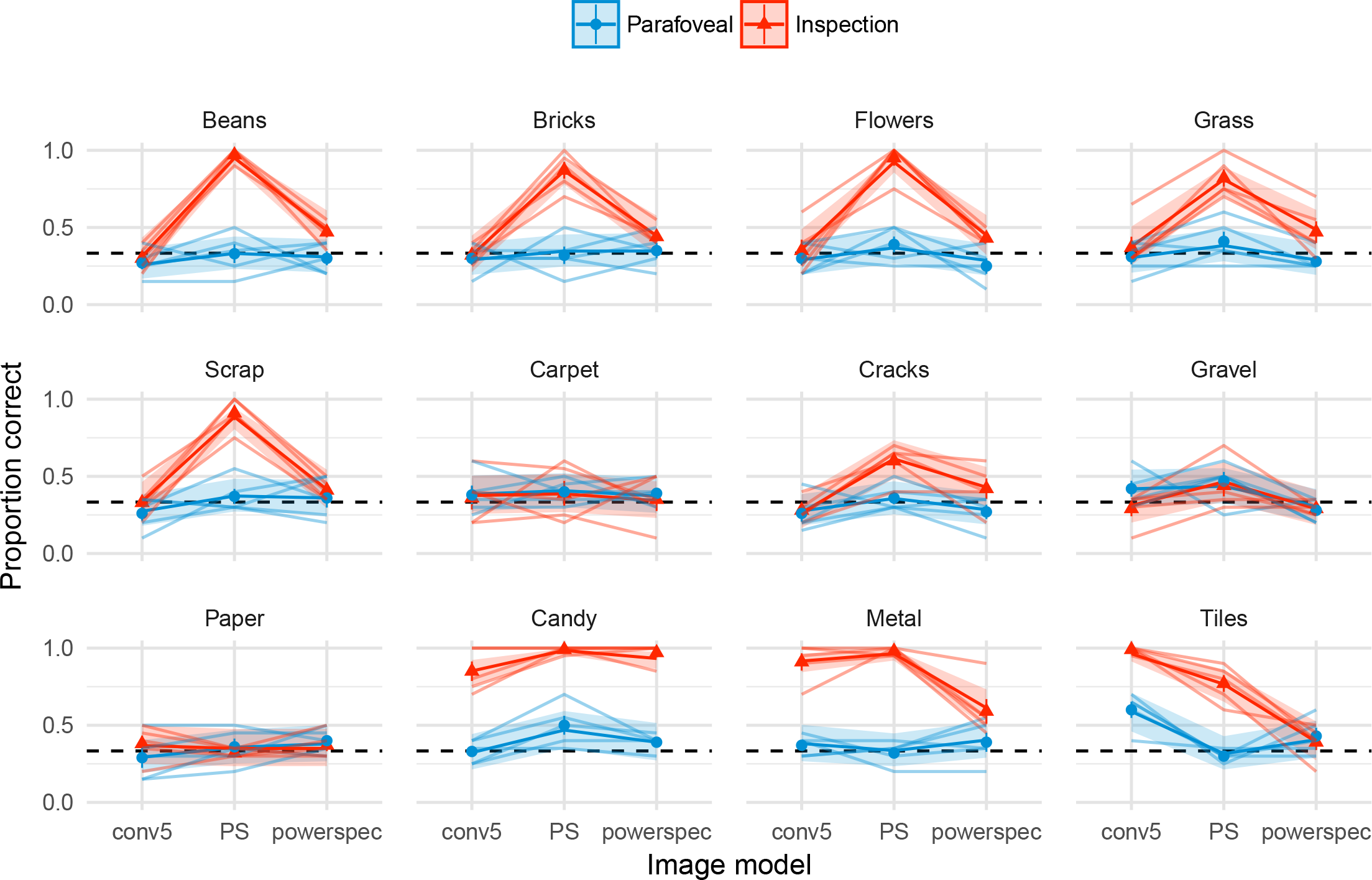
Performance for each image in Experiment 2. Points show means and ± 1*S E M* over observer means. Faint lines link individual observer mean performance (based on 20 trials for all observers except S2). Solid lines show mixed-effects model estimates (posterior mean) and shaded regions show 95% credible intervals.

### Control analysis: cross-correlation of image crops

The experiments reported above show that the CNN texture model (specifically the powerspectrum matching variant) can match the appearance of a range of textures even under foveal viewing. One concern with this result is that the model may be overfitting on the target texture image. Consider a “copy machine” model that would exactly copy the image up to a phase shift. Samples generated by this model would likely be indistinguishable from the original image, because our experimental design (taking non-overlapping crops) enforces the samples to be physically different. Consequently, if a model was acting like a copy machine, this could not show up in our existing results. If this were the case, one could argue that the model has not learned anything about the structure of textures per se but rather how to copy pixels.

To investigate this issue, we computed the normalised maximum cross-correlation between different texture samples and the corresponding original texture. If the algorithm simply copies and phase-shifts the image, the maximum cross-correlation with the original will be one. Specifically, for each of the ten unique texture samples of size 256 × 256 synthesised by each model in Experiment 2, we took one N × N crop of the centre plus ten additional random crops of edge N pixels, for each of N = {32, 64, 128}. Each crop is then normalised to have zero mean and unit variance, before computing the cross-correlation function between crop and original and taking the maximum. Finally, we take the average of this maximum across the eleven crops.

For certain textures however, it may be the case that a synthesis algorithm *needs* to act like a copy machine (up to a spatial shift) to match the appearance of the texture. For example, textures with strong periodicities and little variation between individual texture elements (e.g. Metal or Tiles) might require copying for appearance to be matched, whereas the appearance of less regular structure (Grass, Beans) might be sufficiently captured by far less. To account for this image-specific variation, we additionally computed the maximum cross-correlation between an N × N centre crop from the original texture, and the full 256^
2
^pixel image itself (after excluding shifts of +/- 16 pixels around the centre, which would trivially return one). This value can be seen as a measure of self-similarity.

The maximum cross-correlation values for the images used in this paper are shown in Figure 11. This result shows that crops of synthesized textures are not more similar to the best matching crop in the corresponding original image than are any two crops taken from the original image. Thus, none of the models are simply copying the original images at any of the spatial scales we tested. The Metal and Tiles images are the most self-similar (grey bars) at all scales, and these were also the images for which adding the powerspectrum constraint to the CNN Texture model helped most (compare conv5 and powerspec cross-correlation values).

**Figure 11.**
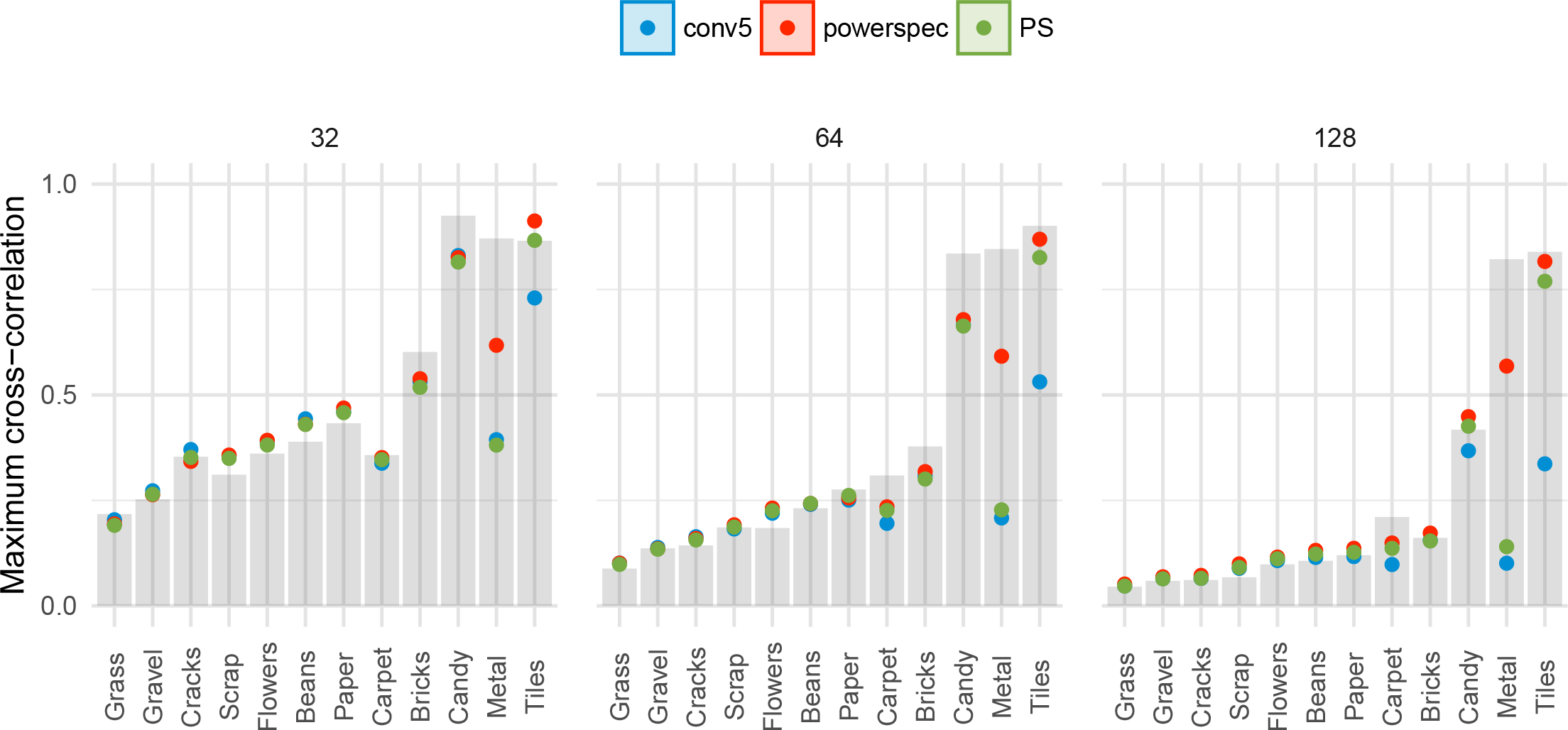
Controlling for texture model overfit. Points show the average maximum cross-correlations between crops of model syntheses (colours) and the original images, for three different crop sizes (32, 64 and 128 pixels; panels). If the model simply copied and phase-shifted the original, these values would be approximately one. Bars show the baseline of a crop from the original image correlated to itself. Some images are more self-similar, and thus require some degree of copying to match appearance.

## General discussion

We have shown that the CNN texture model of Gatys et al. (2015) can produce artificial images that are indisciminable from a range of source textures even under foveal viewing. That is, images synthesised from the Gatys model could pass as natural materials, at least for 9 of the 12 images we test here and for similar viewing conditions. A model that matches both a selection of deep CNN features and the power spectrum of the original image (Liu et al., 2016) greatly improves the perceptual fidelity of two of the remaining three images not captured by the Gatys model (Experiment 2). These results were not attributable to simply copying the target images (Figure 11). The most popular existing par metric texture model (PS; Portilla & Simoncelli, 2000) can capture texture appearance for many images briefly presented to the parafovea, but is less successful under foveal inspection (matching appearance for 4 of the images—see Figure 7). These results regarding the PS model corroborate the findings of Balas (2006) and Balas (2012) respectively. Taken together, our results show that the natural image statistics represented by the CNN model (and the powerspectrum variant) can capture important aspects of material perception in humans, but are not sufficient to capture the appearance of all textures.

The patterns of performance in Figures 7 and 10 suggest that for the purposes of assessing parameteric texture models, natural textures may be parsed into atleast four clusters^
6
^. First, one cluster of images (Beans, Bricks, Flowers, Grass and Scrap) can be matched by the CNN texture model’s higher layers even for foveal inspection, but only for parafoveal viewing by the PS model. These images feature readily-discernable texture elements that do not follow a regular periodic arrangement. The second cluster (Carpet, Cracks, Gravel and Paper) can be matched by all but the simplest CNN texture model under both parafoveal and inspection conditions. For these images, it is possible that individual textons (single texture elements; Julesz, 1981) were difficult to resolve even foveally, allowing models that failed to capture individual textons to nevertheless sufficiently match appearance. Third, the Metal and Tiles images include regular structure that can only be effectively matched by the CNN+powerspectrum model. These are both strongly periodic textures with easily resolvable textons. Finally, the Candy image cannot be matched by any of the models tested here for foveal inspection. It contains large textons with interesting material properties (glossiness^
7
^) as well as occlusions and shading suggesting depth. These clusters may provide useful test cases for parameteric texture models in the future. In particular, a single image from each class may be sufficient to provide a generalisable test of a texture model. More generally, psychophysics may offer an approach to find equivalence classes of textures that are useful for discriminating between texture models^
8
^. The failure of all models we test here to capture the Candy image shows that the CNN features we test here are still not sufficient to capture the appearance of all textures.

An additional noteworthy feature of the data is that for many images, the conv5 model is slightly worse at matching appearance (psychophysical performance is better) than the conv4 model (e.g. Figure 12). This is particularly evident for example for the Candy and Tiles images under inspection (Figure 7, though note these datapoints are also affected by large oddball type biases—see Figure 16). Assuming this effect is robust, it could be related to the observation that the conv5 model results in higher final total loss values after optimization than the conv4 model (Figure 18).

**Figure 12.**
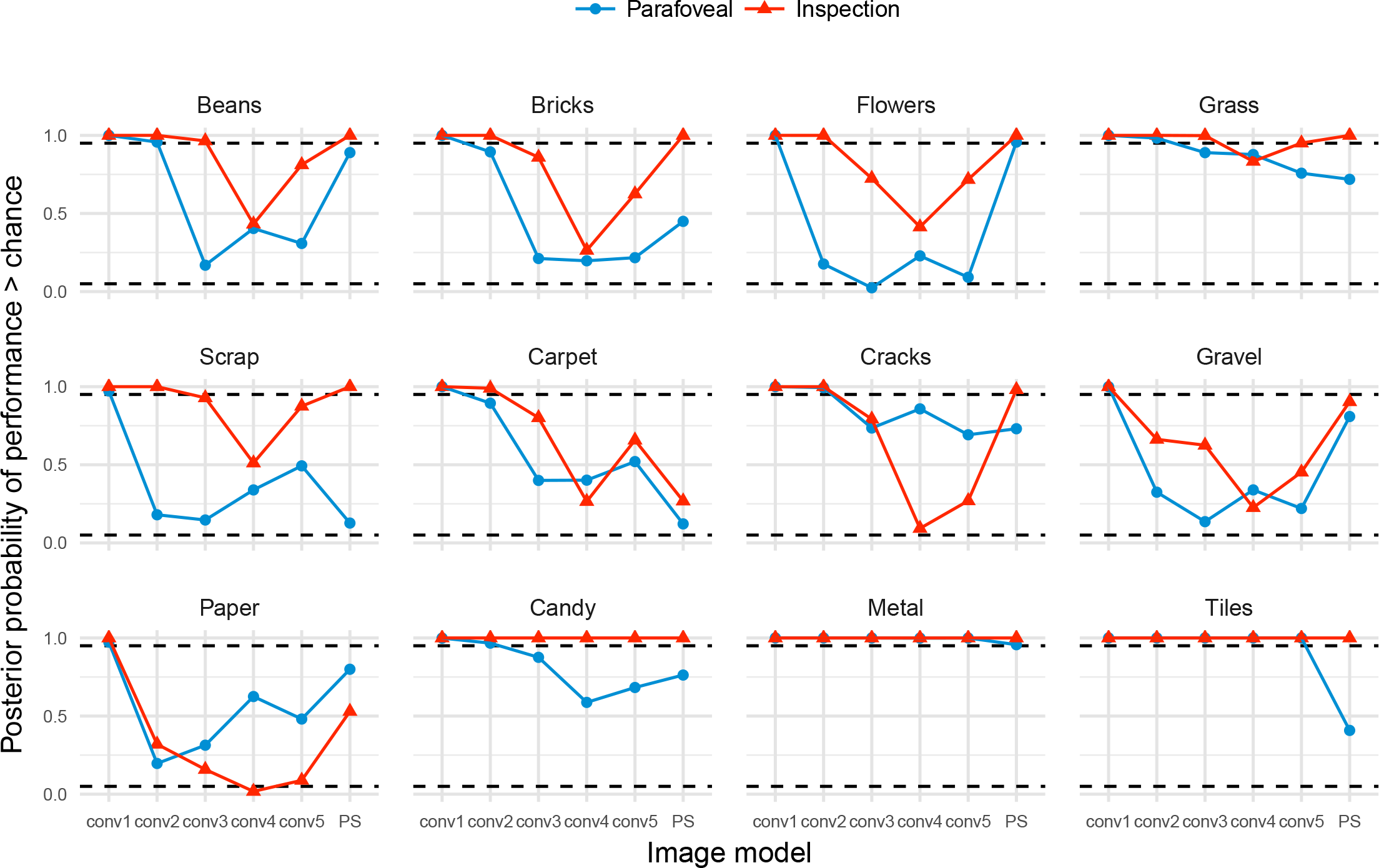
Posterior probability that performance for discriminating each image and image model in Experiment 1 lies above chance. Conditions falling above the dashed horizontal line at 0.95 have a greater than 95% probability of being discriminable, conditions falling below the dashed horizontal line at 0.05 are more than 95% likely to be below-chance. Conditions for which the model predicts exactly chance performance would fall at 0.5.

**Figure 16.**
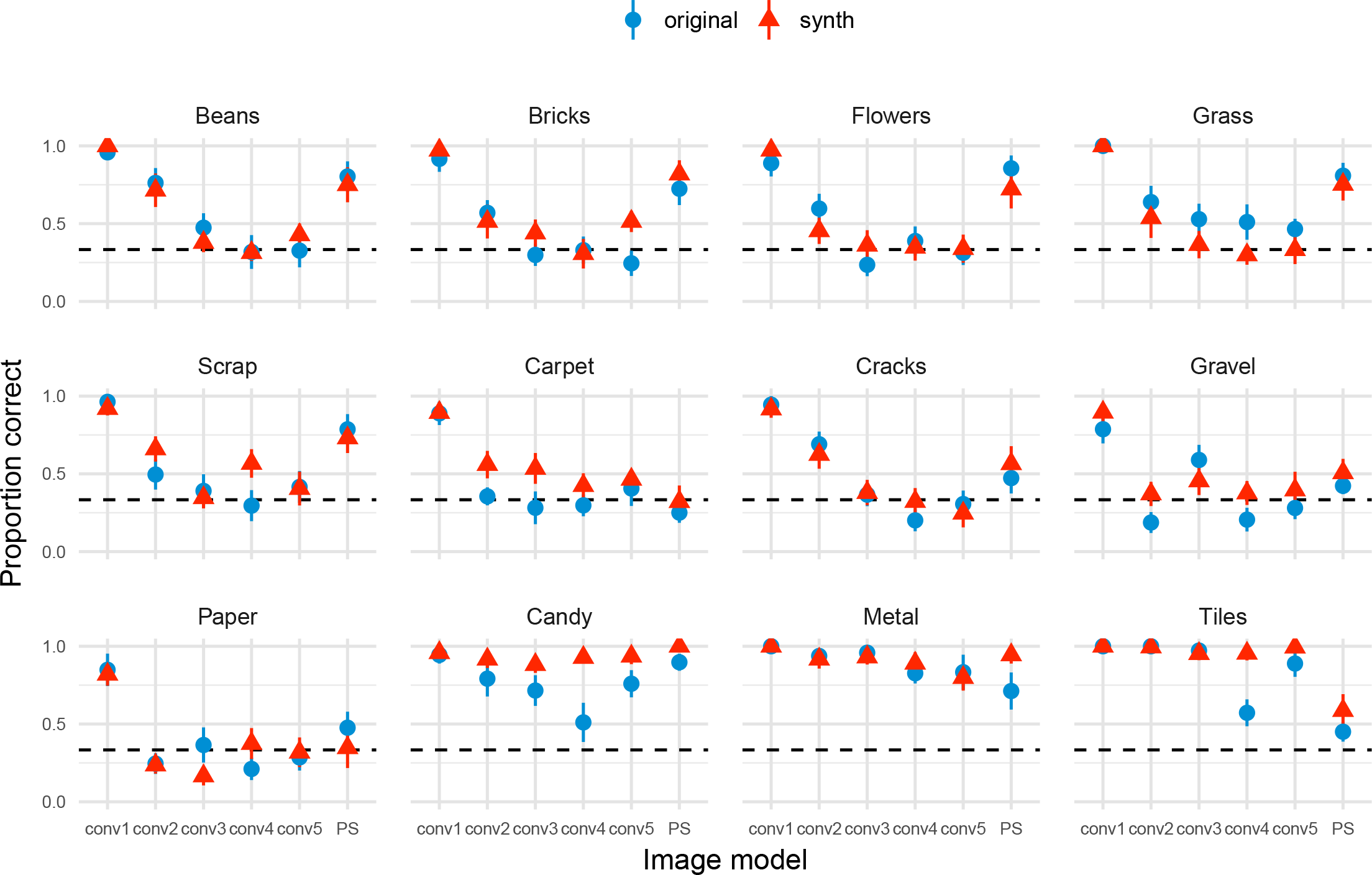
Inspection performance in Experiment 1 according to whether the oddball image was an original or a model synthesis (“synth”), for each image. Points show grand mean across observer means, error bars show SEM.

**Figure 18.**
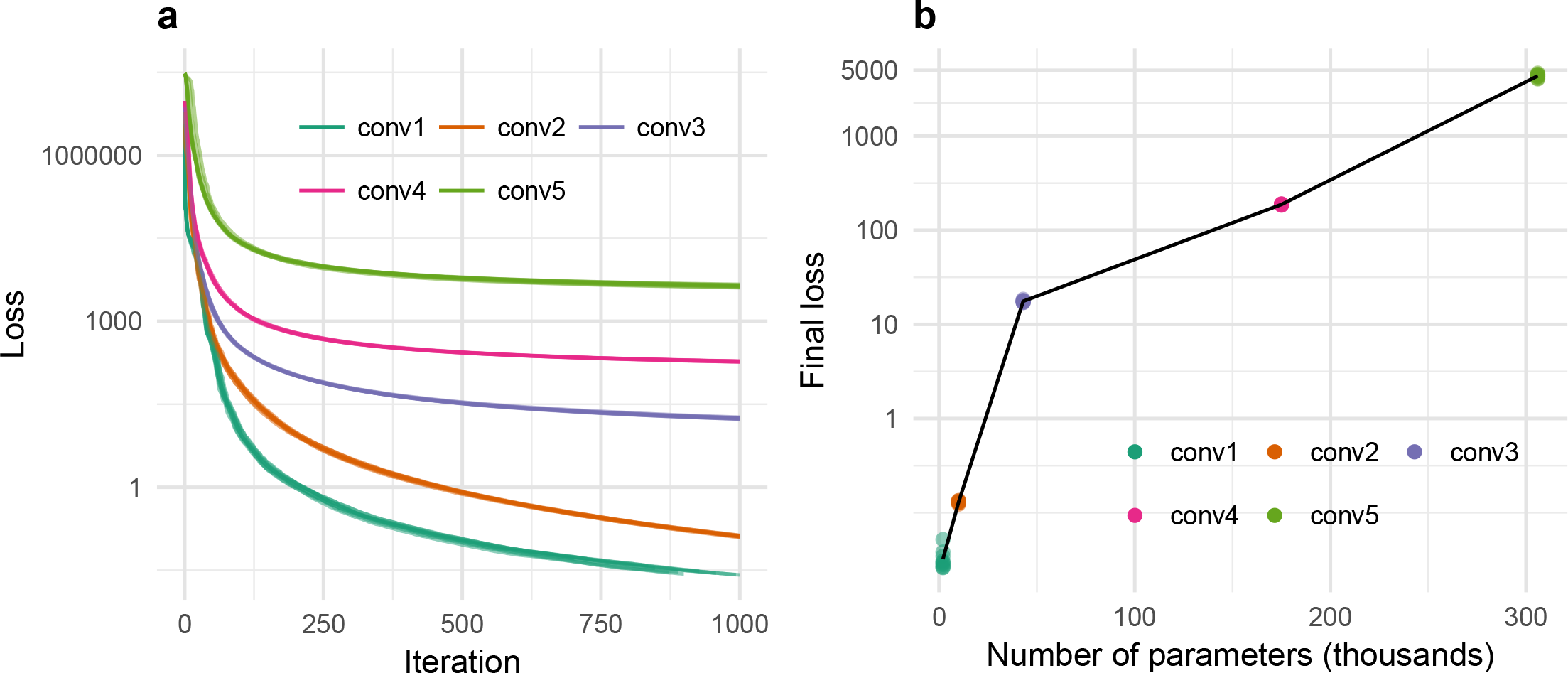
**a**: Decrease of the loss over iterations on a logarithmic scale for ten syntheses (lines) of one example image (Bricks). Loss for simple models (e.g. conv1) converges to zero whereas for more complex models (conv3, conv4 and conv5) it stabilises in a local minimum. **b**: Final loss (logarithmic scale) for the synthesised images in (a) as a function of number of parameters in the model. Points show individual syntheses, lines link means within a model. Final loss is superlinear in the number of parameters.

### Model complexity vs feature complexity

Why do the features used in the CNN texture model often succeed in capturing texture appearance? One possibility is that training to recognise objects in images causes deep CNNs to abstract a set of statistics from images that support quite general visual inferences (transfer learning; Donahue et al., 2013). An alternative possibility is suggested by Ustyuzhaninov, Brendel, Gatys, and Bethge (2016), who found that single-layer CNNs using many filters with random weights could produce textures of surprisingly good perceptual quality (assessed via introspection). That is, high-quality texture synthesis from CNNs may require neither a hierarchical (deep) representation nor filters learned on any particular task—many random filters could instead be sufficient (the Random-Multiscale model from that paper uses about 2 million random parameters, which is significantly more than all models in this paper—Figure 5). If the latter is the case, this would suggest that the improved appearance matching as more convolutional layers are included is because there are simply more features, not that they are “better”.

However, we do not believe the improved appearance matching is only due to the number of parameters matched. Gatys et al. (2015) showed that the number of parameters in the CNN model could be reduced by computing Gram matrices on only the first *k* principle components of each network layer. Textures synthesised using approximately 10,000 parameters from VGG layers conv1_1, conv2_1, conv3_1, conv4_1 and conv5_1 produced (introspectively) much higher-quality textures than only using all parameters from conv1_1 and conv2_1 (about 12,000). A second piece of evidence that speaks to this point is that having more parameters—even having more parameters than pixels (i.e. being overcomplete)—does not necessarily result in introspectively high-quality textures (Ustyuzhaninov et al., 2016). Thus, features from the higher network layers seem to improve texture synthesis because they are “better” features, not simply because they add more parameters.

Why are higher layer network features (with the possible exception of conv5_1; see above) better? Recall that deep convolutional networks stack nonlinearities (Figure 1), allowing increasingly complex functions of the input image to be explicitly (linearly) decoded. Higher layers might therefore be better for texture synthesis because they learn to represent complex information. Alternatively, it could just be that higher layers have larger receptive fields than lower layers, and a model that includes both high and low layer information improves because of its multiscale structure. Ustyuzhaninov et al. (2016) showed that having features at multiple scales improves texture synthesis. On one hand, the fact that trained features produce (introspectively) better textures than the random multiscale network using fewer parameters implies that our texture models including higher VGG layers are not better exclusively because they model information at more spatial scales. Another possibility is that it is easier to optimise trained features than random features, which leads to better texture synthesis but does not mean deep features are “better” for parametric texture modelling in general.

Ultimately we think the models in this paper perform well due to a mixture of both more and more complex features, and that this is not simply a function of including information at multiple scales. Future psychophysical comparisons could be used to add quantitative rigour to this discussion. For example, comparing the perceptual quality of the random-filter and trained CNN model textures (with and without compression) would quantify the importance of learned features. Similarly, comparing hierarchical (cumulative) and non-hierarchical models could be used to quantify the importance of scale information.

Finally, we would like to emphasise that for those textures the CNN model can mimick, the model features likely represent a superset of the necessary statistics. One important challenge now is to compress these representations into a minimal set of features, in order to develop a parsimonious and intuitive description of the critical aspects of the feature space. As noted above, Gatys et al. (2015) showed that qualitatively-reasonable results could be obtained for a PCA-reduced feature space with 10,000 parameters, compared to the 175,000 of the conv4 or 306,000 of the conv5 models used here. Of course, the PS model matches substantially fewer (about 1,000) parameters than even this, and so its performance for parafoveal images is impressive. The difference between the two models, more substantively quantified, could yield insights into the differences in foveal and peripheral encoding of texture.

### Categorical losslessness

Our experiments show that humans cannot tell which of three physically-different images were “generated by a different process” (for all but one of the images we test). This condition could be termed “categorical” or “structural” losslessness (Pappas, 2013): under our experimental conditions, the model syntheses can pass as natural textures (they are perceived as the same category). Images that are perceptually equivalent along some dimension can also be called “eidolons” (Koederink, Valsecchi, van Doorn, Wagemans, & Gegenfurtner, 2017). Achieving categorical losslessness in an image-computable model is an important step towards understanding human material perception, because the model encodes sufficient statistics for capturing the appearance of these textures. Categorical losslessness must however be distinguished from perceptual losslessness: humans are likely able to tell that the three images in our experiments are different from each other (and thus we avoid using the term *metamer* here, which refers to physically different images that cannot be told apart). The latter criterion may be important for understanding information loss in the visual system more generally (Freeman & Simoncelli, 2011; Koenderink & van Doorn, 1996; Wallis et al., 2016; Wandell, 1995).

### Caveats

Three caveats should be borne in mind when interpreting our results. First, we have considered only one relationship between input image size and CNN feature scaling (specifically, we used input images of 256 pixels square, which is close to the 224 pixel square images on which the VGG features were learned). Because the network layers have a fixed receptive field size (the pixels of the original image associated with a given unit), downsampling or upsampling the input images will cause the same network layers to respond to different image structure. For example, it is possible that there is a relationship between the degree to which texture appearance is successfully captured by the model and the size of the texture elements in the image. One possible reason that the Candy image (Figure 2) fails to be matched for foveal viewing is that the textons (individual candies) and their overlap are too large to be captured by single filters at some critical layer within the network, even though features in the highest layers are large enough to cover groups of candies. We have tried rescaling the images but this did not seem to improve the syntheses, indicating that this relationship is perhaps not trivial.

A second caveat is that the fidelity of the resulting textures could depend on the number of iterations of the gradient descent used to minimise the loss between the original and the new image (see Appendix, Figure 18). Because this loss is never exactly zero for the more complex models, more iterations could only improve synthesis fidelity—though in our experience, the coarse structure of the images is largely fixed within 200 iterations, and further iterations mostly reduce high-spatial frequency noise. In theory, as long as all features are perfectly matched (i.e. if the loss is exactly zero), more features can only lead to more similar patterns. However, given that the optimization of texture synthesis algorithms typically yields a residual loss, more features do not necessarily improve perceptual quality, and the design of good features is not straightforward and may depend on various factors including the type of textures to be synthesized. As it stands, different models are ideal for different purposes. For peripheral texture perception the PS model achieves best performance with relatively small number of parameters, for random fields with pairwise interactions the scattering network provides a very compact representation for texture synthesis (Joan Bruna, personal communication) and for foveal inspection of textures the VGG features seem particularly useful.

Finally, in our experiments we closely followed the oddity method used by Balas (2006). We believe this paradigm has many desirable properties as a measure of categorical losslessness, but our results also point to a caveat. By cropping from inhomogenous images (e.g. the Flowers image, which contains a size gradient) we introduce greater perceptual variability in the stimuli shown to subjects. Depending on the relative (in)homogeneity of original and synthesised images, this may lead to differences in performance depending on the class of the oddball and potentially to below-chance performance (e.g. in the Flowers image). We discuss these issues and present further analysis in the Appendix. While we believe this property will have little effect on our overall conclusions, it is nevertheless useful to consider for future studies.

## Conclusion

We have shown that the texture model of Gatys et al. (2015), which uses the features learned by a convolutional neural network trained to recognise objects in images, provides a high-fidelity model of texture appearance for many textures even in the fovea. Overall however, our results do not identify a uniformly best parametric model for matching texture appearance. Instead, different models may be appropriate for different use cases. The PS model is the best (and the most simple) model to use if textures are intended to be viewed briefly in the parafovea. For textures intended to be foveated, incorporating the powerspectrum constraint will be critical for textures with strong periodicities (Liu et al., 2016), whereas the CNN model (conv4) performs best for most other textures we test here. It would obviously be desirable to identify a uniformly best model in future work, and the single failure case we identify here (the “Candy” image) may provide a useful benchmark for testing such models.

## Acknowledgements

### Acknowledgments

Designed the experiments: TSAW, ASE, CMF, LAG, FAW, MB. Programmed the experiments: TSAW. Collected the data: CMF, TSAW. Analysed the data: TSAW, AE, CMF. Wrote the paper: TSAW. Revised the paper: CMF, ASE, LAG, FAW, MB. Thanks to Paul-Christian Bürkner for his assistance with fitting the GLMMs in brms, Heiko Schütt for helpful comments on presentation, and www.textures.com for permission to use the images. Funded, in part, by the German Federal Ministry of Education and Research (BMBF) through the Bernstein Computational Neuroscience Program Tübingen (FKZ: 01GQ1002), the German Excellency Initiative through the Centre for Integrative Neuroscience Tübingen (EXC307), and the German Science Foundation (DFG; priority program 1527, BE 3848/2-1 and Collaborative Research Centre 1233).

## Appendix

### Bayesian multilevel modelling

To analyse the data, we first made the (standard) assumptions that the observers’ responses on each trial (correct / incorrect) reflected a Bernoulli process, and that the response on a given trial was not dependent on previous responses. We estimate the success probability of this Bernoulli process using a generalised linear mixed-effects model (GLMM) with a logistic link function whose parameters were estimated in a Bayesian framework^
9
^. A mixed-effects model (a type of hierarchical or multilevel model) includes some number of “fixed” effect parameters that quantify how the response depends on the independent variables at a population level, and some “random” (also called “group-level”) effects that allow the fixed effect coefficients to depend on discrete levels that are assumed to be non-exhaustive samples from a larger population. Our model contains two fixed-effect factors: the image model (with six levels, entered into the model design matrix using successive difference coding using contr.sdif from the MASS package for R; Venables and Ripley (2002)) and the presentation condition (with two levels, parafoveal and inspection, coded with sum contrasts [1, -1]). We included the interaction terms between these factors such that the model consisted of 12 fixed effect coefficients. The variation caused by observers and images are modelled as random effects, which are coded as offsets added to the fixed effect coefficients whose variance is estimated. Note that we make an additional simplifying assumption by ignoring other sources of variance, such as the synthesised image used on a trial and the random crop location (see Methods). We assume that each fixed effect coefficient can vary by observer and / or by image, and that the variance could be correlated. The specification of the model in R formula syntax (lme4 / brms) was

> `model_formula <- correct ~ image_model * presentation_cond +`
>
> `(image_model * presentation_cond | subj) + (image_model * presentation_cond | image_code)`

We used conservative, weakly-informative prior distributions in the sense that they bias estimates towards the middle of the range of possible values and away from indicating large effects. Consider that the model coefficients are defined on the linear predictor scale, whose effective range runs from approximately -5 (returning an expected success probability of 0.007) to 5 (returning 0.993; a linear predictor value of zero gives 0.5). We therefore expect that no standardised fixed-effect coefficient to be larger than ± 5 (i.e. the difference between two factor levels runs from the lowest to the highest observable success probabilities, other effects being equal), and they will very likely be smaller than this. We therefore place Gaussian priors over all fixed-effect coefficients for factors with mean zero (i.e. our *a priori* expectation is for no effect), standard deviation 2 (indicating a weak implausibility of large coefficients). These are therefore weak, but not flat (uniform) prior distributions. We also place priors over the variances for random effects; following the logic for effective range of the linear predictor we expect that the effect sizes of our fixed effects are unlikely to vary by more than 2 on average (i.e. the standard deviation is very unlikely to be larger than 2). We use half-Cauchy priors (i.e. with a lower-bound of zero, as recommended by Gelman & Hill, 2007) over the standard deviation parameters for each random effect, with a mean of zero (i.e. our maximum a priori assumption is that subjects and images are no different) and a standard deviation of 1, reflecting large uncertainty. Finally, we set a prior over the correlation matrix for observer and image-level offsets in the fixed effects that assumes that smaller correlations are slightly more likely than larger ones (an “lkj(2)” prior, see Lewandowski, Kurowicka, and Joe (2009); Stan Development Team (2015) for details). While the priors we use here are informed by the scale of the model and by common practice for Bayesian regression models (see for example Gelman, 2006; Gelman & Hill, 2007; Kruschke, 2011), the specific choices we make are somewhat arbitrary. As we see above, the model provides a good fit to the data, but the reader should bear in mind that as always, our inferences depend on the model we assume.

We estimate the posterior distribution over model parameters using a Markov Chain Monte Carlo (MCMC) procedure implemented in the Stan language (version 2.15.1; Hoffman & Gelman, 2014; Stan Development Team, 2017), using the model wrapper package brms (version 1.7.0; Bürkner, in press) in the R statistical environment. The brms package allows the specification of flexible mixed-effects Stan models using formula syntax similar to the popular lme4 package (Bates, Mächler, Bolker, & Walker, 2015). Samples were drawn using the NUTS sampling algorithm (Hoffman & Gelman, 2014) with 6 independent chains, each sampled with 30000 samples of which 10000 were used to adaptively tune the sampler (warmup). To reduce the final file size we saved every 6th sample. This procedure resulted in a final total of 20000 post-warmup samples. Chain convergence was assessed using the Rˆ statistic (Gelman & Rubin, 1992) and visual inspection of traceplots. Readers are encouraged to consult the online code for further details.

The resulting posterior distribution is summarised as Bayesian credible intervals on marginal parameter values and predictions. Unlike frequentist confidence intervals in general, credible intervals have the desirable property that they represent a coherent statement of belief about the parameters’ likely values, given the model, priors and data. A 95% credible interval means that the “true” parameter value (conditioned on model, prior and data) has a 95% probability of lying within the interval (see Miller & Ulrich, 2015; Morey, Hoekstra, Rouder, Lee, & Wagenmakers, 2015; Morey, Hoekstra, Rouder, & Wagenmakers, 2015, for recent discussion on this issue), which many readers will find intuitively appealing. We report 95% credible intervals (rather than 67% or 82% or any other interval) merely as convention. The model’s belief about the data is represented by the full posterior distribution, which can be summarised into arbitrary intervals (see McElreath, 2016, p.58 for related discussion). Readers should avoid mental hypothesis testing (rejecting null values that lie outside the interval). Using Bayesian credible intervals to reject null values in this way suffers two of the same problems as null hypothesis significance testing using p-values: it can only reject but never accept a null value, and if used with optional stopping of data collection it will always reject null values even if they are true (Kruschke & Liddell, 2017). Instead, the credible intervals serve to give information about the magnitude and precision of likely effects.

Another advantage of a Bayesian approach in this context is that the weakly-informed priors we use act as a regulariser for the model, ensuring that parameters are identifiable (indeed, in our hands the lme4 package had troubles fitting this model). Using zero-centred prior distributions on regression parameters biases the parameters against finding spuriously large effects. One caveat is that credible intervals in general, unlike confidence intervals, are not guaranteed to result in a pre-specified error rate for binary inferences (e.g. effect / no effect) in the long run. Given that some decisions about our analyses were made after seeing the data (making this *exploratory* research), frequentist p-values would not have their nominal false-alarm rates in any case. For these reasons we report a Bayesian analysis here; readers wishing to apply other analyses are encouraged to do so using the raw data provided online.

Where it makes sense to compare discrete models, we do so using an approximation to the out-of-sample (leave-one-out) prediction error provided by the R package loo (v 0.1.6; Vehtari, Gelman, & Gabry, 2016). Loosely, this value estimates the ability of the model to predict new data (smaller values are better). We report differences between models and their standard errors on the deviance scale (2 the expected log point wise predictive density estimated by the loo package, called LOOIC).

**Experiment 1.** Figure 6 shows model predictions for both individual observers (authors CF and TW) and for the average of the naïve observers. For the individual observer model estimates (CF and TW) we show the model prediction conditioned on observer. The observer’s mean performance is 95% likely to lie within the shaded area for an average, unknown image (Baayen, Davidson, & Bates, 2008). The “naïve” panel shows the average performance for the naïve observers. The model predictions here exclude both observer and image random effects: mean performance has a 95% probability to lie within the shaded area for an average, unknown image and an average, unknown subject. Note that the model uncertainties shown in Figure 6 depict the expected spread of population averages across images, but are not appropriate for comparing between presentation conditions because they do not take into account the paired nature of these data (the design was within-subjects and within-images).

To quantify the differences between conditions more appropriately we examine the mixed-effects model coefficients. First, we quantify the performance difference between the inspection and parafoveal conditions, marginalising over image models and all random effects variance. The posterior median of the difference between these conditions on the linear predictor scale is 1.23. Considering the exponent of this value as log odds, this means that correct trials are exp(1.23) = 3.41 times more likely under the inspection condition than the parafoveal condition, if all other effects are held at zero. In other words, for every 10 correct responses in the parafoveal condition we expect about 34 correct responses in the inspection condition, on average. The 95% credible interval tells us to believe that the difference has a 95% probability (conditioned on the data, model and prior) of lying between 0.65 and 1.82. To indicate the likely sign of an effect we report the posterior probability that the coefficient is negative (if this value is small, the coefficient is likely positive; if the value is 0.5 then the coefficient is equally likely to be positive or negative). The inspection condition is very likely to elicit higher performance than the parafoveal condition, because the coefficient coding their difference has only a small probability of being negative (p(*β* < 0) = 9.998e-05). To make future quantifications more concise, for the remainder of this section we report them as (*β* = 1.23, 95% CI = [0.65, 1.82], p(*β* < 0) < 0.001).

Next, we examine whether the differences between image models depended on the presentation condition. An interaction is clearly evident in Figure 6. This subjective impression was supported by a model comparison between a linear and an interaction model using a measure of each model’s ability to generalise to new data (the LOOIC; the interaction model had a lower LOOIC by 294 (SE = 33)). We therefore further consider the differences between image models conditioned on the presentation condition.

For the parafoveal condition, image models above conv2 and also the PS model produced performance at approximately chance level (see below). Our model quantifies the sequential differences between the models, with the coefficients coding the difference between two models on the linear predictor scale. Performance in conv2 was worse than conv1 (*β* = -1.35, 95% CI = [-1.94, -0.78], p(*β* < 0) > 0.999), and conv3 was worse than conv2 (**β** = -0.26, 95% CI = [-0.58, 0.06], p(*β* < 0) = 0.947). However, because performance was now approximately at chance, there was no evidence that conv4 was different to conv3 (β = 0, 95% CI = [-0.26, 0.29], p(*β* < 0) = 0.487) or that conv5 was different to conv4 (*β* = -0.02, 95% CI = [-0.25, 0.21], p(*β* < 0) = 0.561). Similarly, the PS model was also not different to conv5 (*β* = -0.01, 95% CI = [-0.51, 0.52], p(*β* < 0) = 0.508).

The inspection condition showed similar results as the parafoveal condition with two exceptions: first, performance remained approximately above chance, and psychophysical performance was better for the PS model than the conv5 model (i.e. synthesised and natural textures were easier to discriminate). The conv2 model produced worse performance than conv1 (*β* = - 3.08, 95% CI = [-3.87, -2.32], p(**β** < 0) > 0.999) and conv3 produced worse performance than conv2 (**β** = -0.62, 95% CI = [-0.98, -0.26], p(**β** < 0) = 0.999). Conv4 produced worse performance than conv3 in that the coefficient coding their difference was likely to be negative (*β* = -0.38, 95% CI = [-0.68, -0.09], p(*β* < 0) = 0.995). Performance for the conv5 model was approximately equal to conv4 (*β* = 0.19, 95% CI = [-0.05, 0.43], p(*β* < 0) = 0.056). Finally, there was weak evidence that PS model produced better psychophysical performance than the conv5 model when observers could inspect the images (*β* = 0.82, 95% CI = [0.09, 1.54], p(*β* < 0) = 0.014).

To summarise, the two most important characteristics of these data are first, that psychophysical performance is effectively at chance for the parafoveal condition for the conv4, conv5 and PS models. Second, under inspection the PS model produces poorer matches to appearance (better psychophysical performance) than the conv5 and conv4 CNN texture models. Taken together, the data show that the PS model features are sufficient to capture the appearance of natural textures under brief, parafoveal viewing conditions, but that the increased complexity of the CNN model features improves appearance-matching performance under inspection.

The attentive reader may wonder why the model’s uncertainty estimates in Figure 6 are so large relative to the confidence intervals on the data (particularly in the author plots, which are quite precisely measured). We believe this highlights a particular strength of mixed modelling for psychophysical data (Cheung, Kallie, Legge, & Cheong, 2008; Knoblauch & Maloney, 2012; Moscatelli, Mezzetti, & Lacquaniti, 2012): multiple sources of variability can be accounted for and incorporated into predictions at various levels (e.g. the observer and image level, or the subject level ignoring images, or the population level). In this case, averaging over the images and displaying credible intervals that ignore the pairwise experimental design (as in Figure 6) disguises the fact that different images show distinctly different effects of image model and presentation time. For example, for each fixed effect coefficient we can ask whether more variance in the data is caused by variation over observers or images. On average, the variance associated with images is 2.1 times greater than that associated with observers. The linear predictor difference between PS and conv5 averaged over presentation condition is associated with about 3.3 times more variance from images than from observers. That is, this difference tends to depend strongly on the image (Figure 7). The model uncertainties in Figure 6 are large because the “average” or population-level behaviour is uncertain in light of this; indeed, it may make little sense to talk about a “population-level” over images from these data. In contrast, Figure 7 shows model estimates that are far more constrained relative to Figure 6, because the uncertainty in the estimates now reflects between-subject variability rather than between-image variability.

Chance performance in the oddity task indicates the original and synthesised images are not discriminable from each other. To what degree do our data suggest observers perform above chance for each image and viewing condition? One way to quantify this is to compute the proportion of posterior probability density lying above chance performance. This estimates, for every condition, the probability of observers being sensitive to the difference between original and synthesised textures. Conditions that lie above the dashed horizontal line are those for which we can be more than 95% certain (conditional on model and priors) that observers are sensitive to the difference between original and synthesised images. These dashed lines are provided as a guide rather than to encourage dichotomous decision making about “different or not”. The posterior probabilities confirm, in general, our qualitative statements made in the manuscript (Figure 12).

**Experiment 2.** The results of Experiment 2 for the conv5 and PS models replicate the results of Experiment 1. When stimuli are presented briefly to the parafovea, observers are effectively at chance to discriminate both conv5 and PS from the original textures, and there was evidence that the models did not differ (*β* = 0.15, 95% CI = [-0.4, 0.71], p(*β* < 0) = 0.275), whereas under inspection the PS model was easier to discriminate from the original images than the conv5 model (*β* = 1.45, 95% CI = [0.48, 2.41], p(**β** <0) = 0.003). Additionally matching the powerspectrum (“powerspec” model) produced similarly indistinguishable performance from the PS model in the parafovea (*β* = -0.13, 95% CI = [-0.63, 0.36], p(*β* < 0) = 0.712), but better performance than the PS model under inspection (*β* = -1.64, 95% CI = [-2.35, -0.95], p(*β* < 0) > 0.999).

Posterior probabilities that performance lies above chance for each image and viewing condition are shown in Figure 13. As for Experiment 1, these values generally support our qualitative statements made in the manuscript.

**Figure 13.**
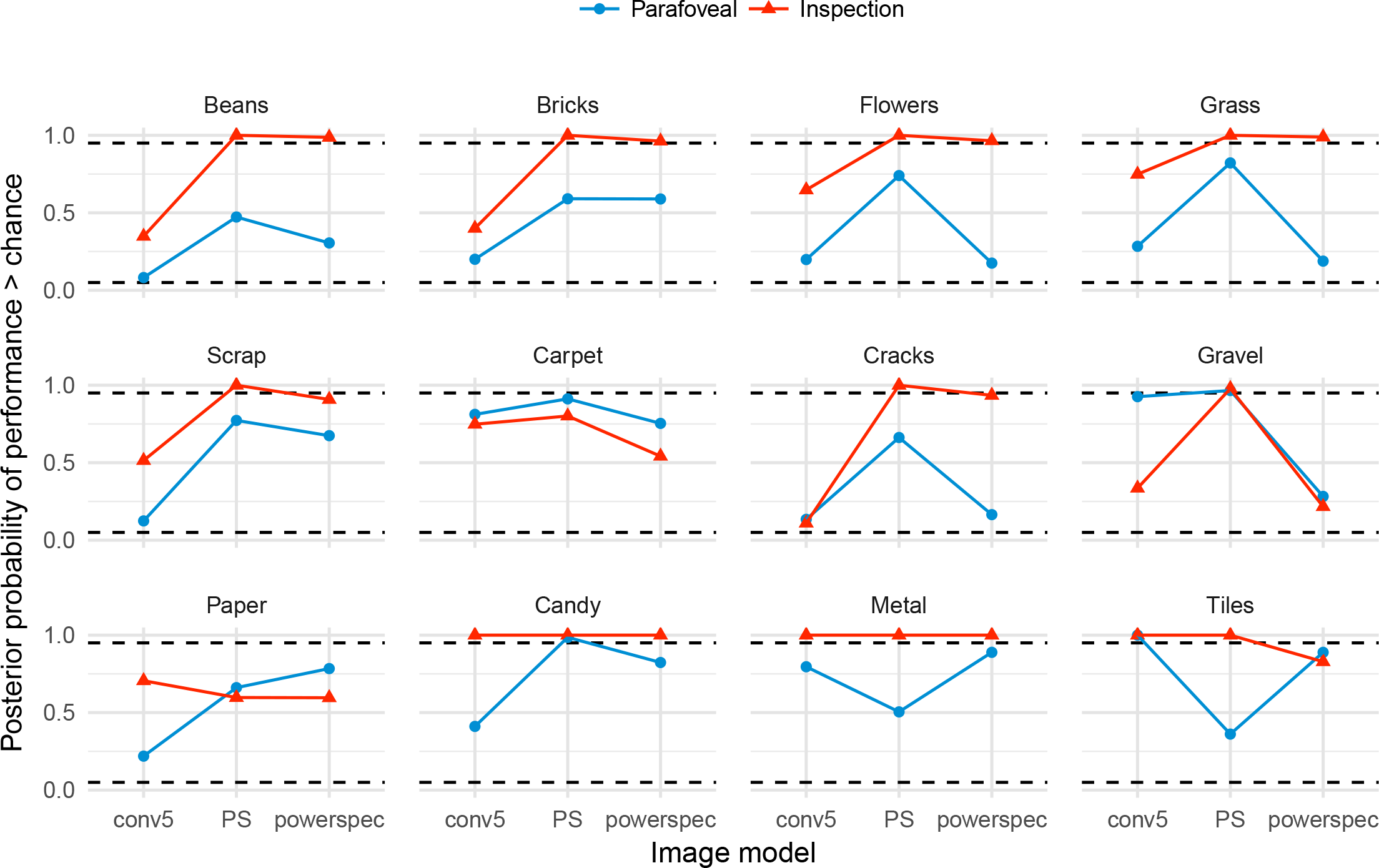
Posterior probability that performance for each image in Experiment 2 lies above chance. Plot elements as in Figure 12.

### Performance as a function of oddball type

Consider that some data points appear to be reliably *below* chance performance (see for example the conv3 model in the Flowers image). Below-chance performance in a forced-choice task generally only occurs in observed data due to measurement error or to observers incorrectly switching responses. However, in our experiments, it is also possible that below-chance performance could be caused in part by cropping from inhomogenous images. For example, the original Flowers image (Figure 2) contains a size gradient such that flowers on the bottom are larger and more sparse than flowers on the top of the image, and this size gradient may result in greater inhomogeneity in the synthesised textures. More generally it may be the case that performance will depend on the relative (in)homogeneity of the original or synthesised images.

To investigate this further we computed performance for trials where the oddball image was an original compared to a model synthesis. When averaging over observers and images (Figure 14), performance is generally slightly higher if the oddball image is a model synthesis rather than an original image. The size of this effect depends on the particular image. For example, in the parafoveal viewing condition (Figure 15) the advantage for synthetic oddballs is quite strong for Metal and Tiles. Similarly, under inspection (Figure 16) observers remain highly sensitive to oddball Candy and Tile syntheses, whereas their performance is relatively poor when the oddball is an original image. This seems particularly strong for the conv4 model, explaining the lower average performance under this model condition.

**Figure 14.**
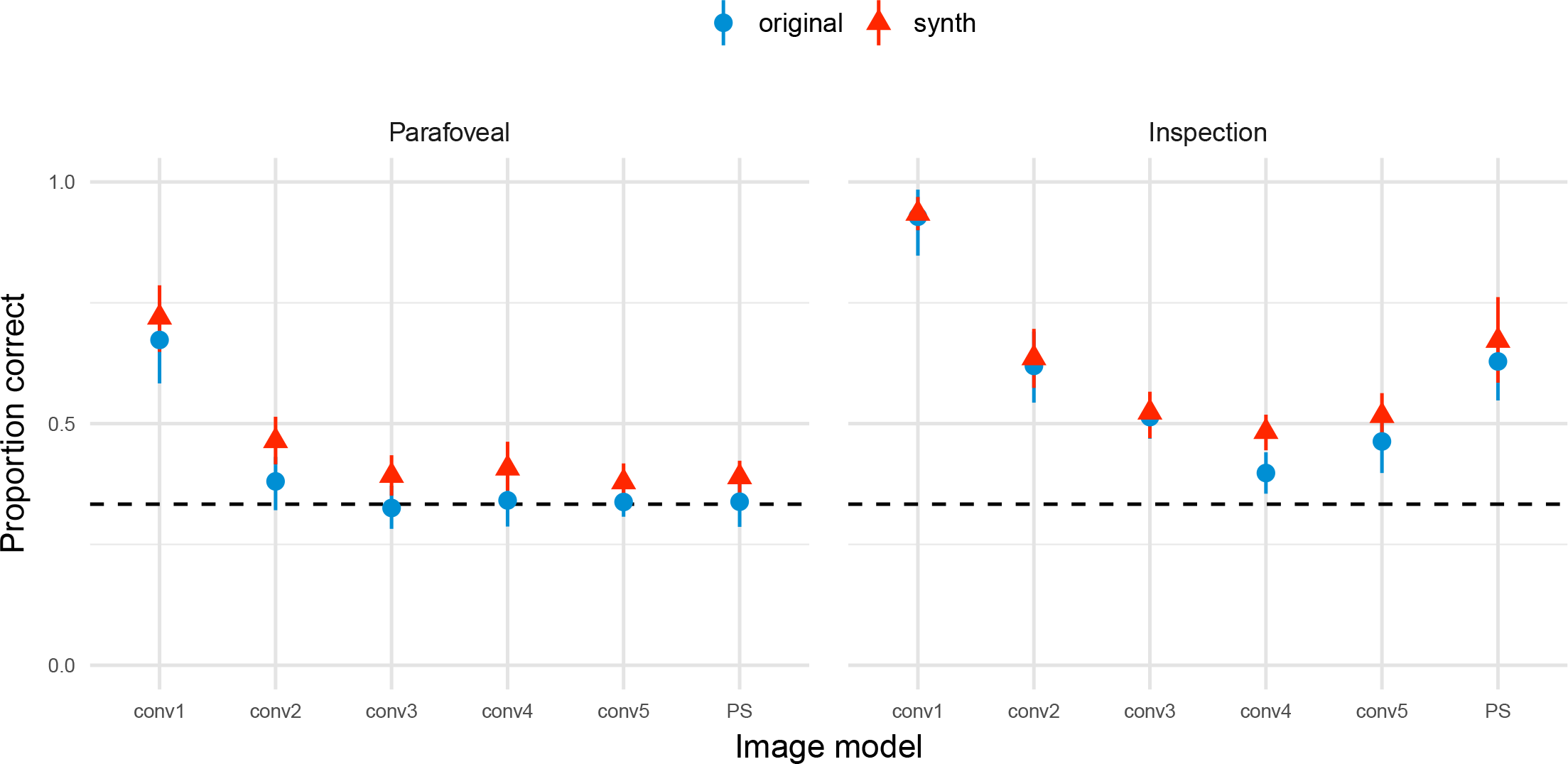
Performance in Experiment 1 according to whether the oddball image was an original or a model synthesis (“synth”), averaging over images. Points show grand mean across observer means, error bars show SEM.

**Figure 15.**
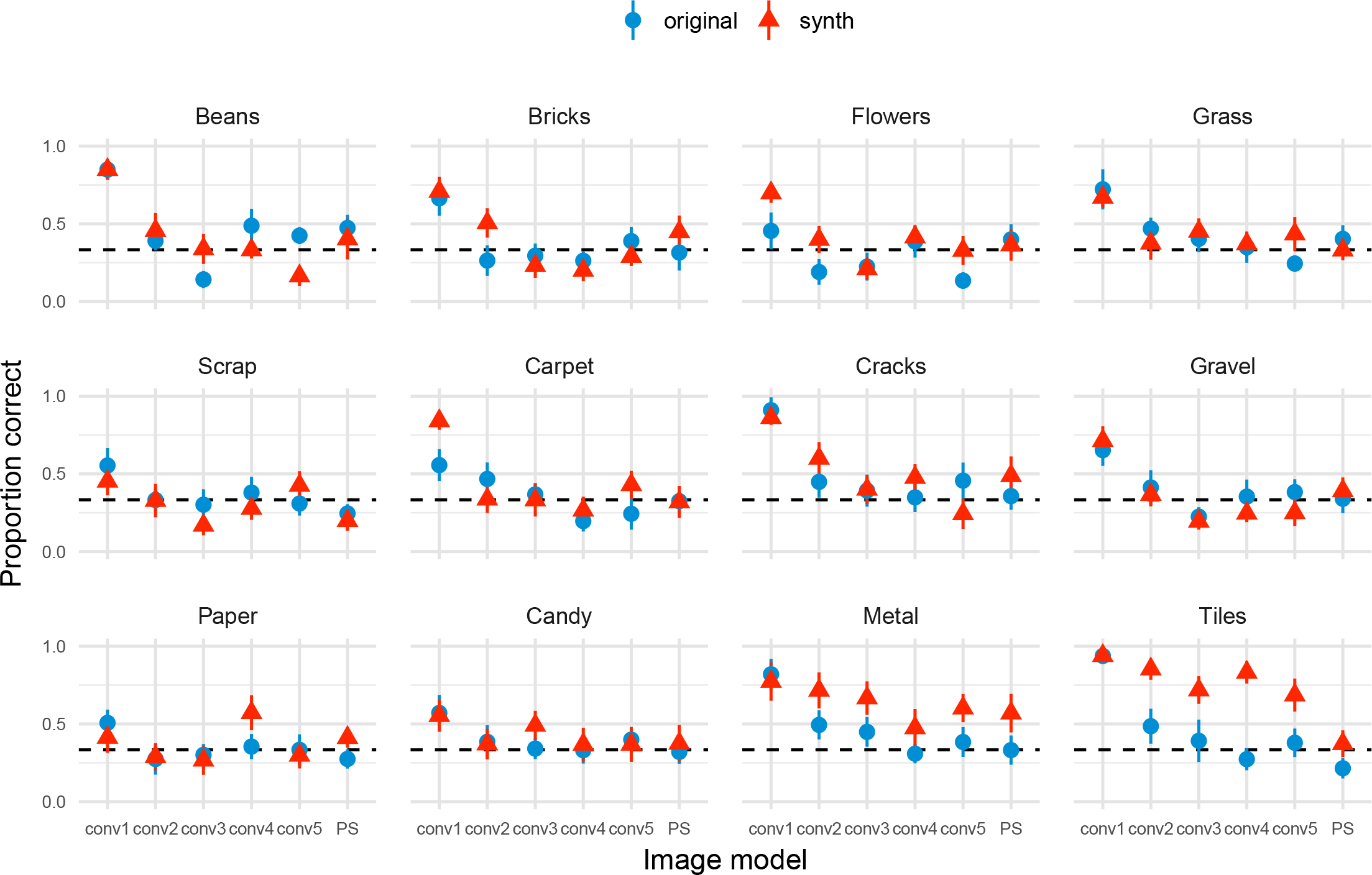
Parafoveal performance in Experiment 1 according to whether the oddball image was an original or a model synthesis (“synth”), for each image. Points show grand mean across observer means, error bars show SEM.

These differences according to oddball type are generally consistent with the perceptual variability account above. If crops from the synthesised images appear different to each other and to the original, but crops from the original are quite self-similar, then on trials with an original oddball each of the three images looks different to the others. One of the synthesised images may appear “most different” (Figure 17a), and the observer incorrectly chooses that. Conversely, on trials where the synthesised image is the oddball, the two intervals containing the original images look similar to each other but different to the synthesised image (Figure 17b), making the task easier. This perceptual variability explanation is particularly appealing for images where the model fails to match appearance, such as for Candy, Metal and Tiles, and is also consistent with the larger self-similarity of those images (Figure 11). Other, not mutually-exclusive, possibilities include that observers are influenced by non-perceptual factors, such as the use of a sub-optimal decision strategy (“pick the unnatural-looking image”) on some trials, or of exogenous orienting of spatial attention to unnatural images. Whatever the cause(s) of the oddball differences we observe, note that traditional observer models for the oddity paradigm assume both unbiased responding and that the stimulus classes have equal variance (Macmillan & Creelman, 2005, p. 235); thus, computing *d*
^′^from our data with the intention of comparing sensitivity to other paradigms should be performed cautiously or with a model explicitly including bias / variance terms for each trial type.

**Figure 17.**
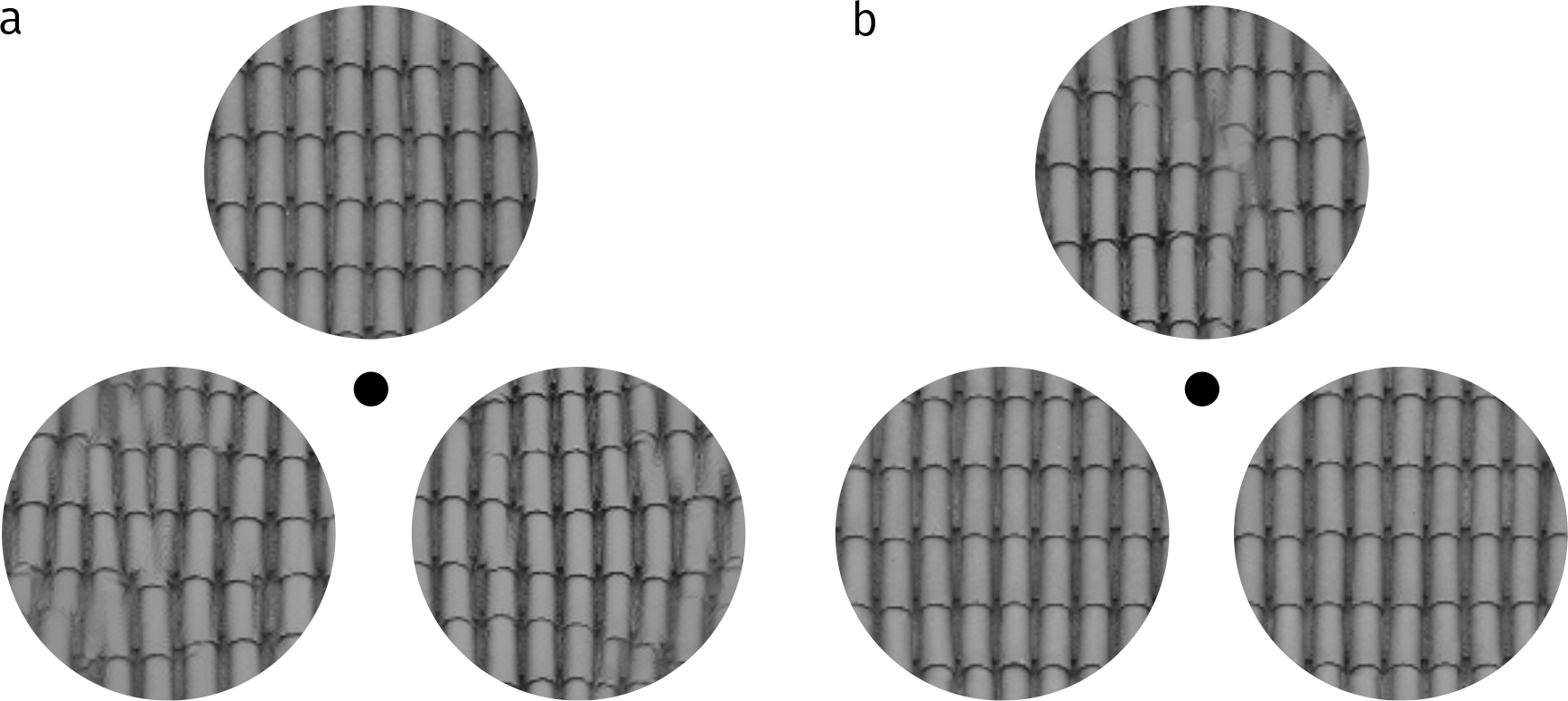
A depiction of an oddball “original” trial (a) and an oddball “synth” trial (b). In both cases the oddball is the top image. All images are physically different. When model syntheses look different to the original and each other, and the original images are very self-similar, then the perceptual variability of all stimulus intervals is larger on oddball original than oddball synth trials.

### Loss

For the stimuli used in this study, the CNN texture models conv4, conv5 and powerspec are overcomplete (have more parameters than pixels in the image). Thus the loss of the gradient descent for those models does not converge to zero, but ends in a local minimum. Figure 18A shows a typical convergence function, where the gradient descent for conv1 terminates early (after reaching convergence within tolerance) but for more complex models (conv3–conv5) loss appears to find a local minimum, remaining relatively stable after 750 iterations. The final loss after 1000 iterations is super-linear (Figure 18B): for example, conv5 has a little less than double the number of parmameters as conv4, but about 23 times higher final loss.

Given that we interleaved ten unique syntheses for each original image within our experiment, it would be interesting to assess whether a correlation exists between the final loss of each synthesis and psychophysical performance. A positive correlation between loss and performance would mean that images that show greater difference to the original under the model would also be easier for humans to discriminate. Unfortunately however, we did not save the final loss of the images after gradient descent but prior to histogram matching. Because histogram matching substantially alters the loss values under the model, including changing the order of syntheses, we are unable to assess a correlation between performance and final loss in this dataset.

## Footnotes

1 As Balas (2006) writes, “the 3AFC task presented here represents a modest contribution towards the formulation of texture discrimination tasks that make explicit the importance of local texture analysis in the human visual system.” We agree.

2 These are analogous to Balas’ “preattentive” and “attentive” conditions, but we consider these terms somewhat of a historical misnomer: because there is no spatial or temporal uncertainty, observers can presumably accurately deploy spatial attention to the stimuli in both cases.

3 These images are copyrighted by www.textures.com (used with permission). Copies of the texture images used in the experiments are available with the online materials of this article (redistributed with permission).

4 Ten was chosen *a priori* based on pilot testing.

5 Observer S9 completed 144 trials of the "Inspection" condition before this data was lost due to computer malfunction. The observer repeated the full testing session; thus this observer had more practice and exposure to the images than the other observers.

6 Since we have only used 12 texture images in the present study, it is likely that a number of additional clusters exist that were not represented in the set of images we used.

7 While the structure of the Candy image is never successfully captured by the CNN model, one intruiging feature of the syntheses is that they appear glossy as for the original image (compare for example the conv3 and conv4 syntheses in Figure 3). This glossy appearance is not captured by the PS model.

8 Balas (2006) subjectively delineated three texture categories: *pseudoperiodic* (containing strongly periodic structure), *structured* (repeated structural elements with no periodicity) and *asymmetric* (containing asymmetric lighting giving the impression of depth). Our cluster containing Metal and Tiles is equivalent to Balas’ pseudoperiodic textures, but our other three data-determined clusters do not trivially map onto Balas’ other categories (e.g. Bricks and Grass are structured, whereas Flowers, Beans and Scrap contain asymmetric lighting and other depth cues).

9 A three-alternative forced-choice procedure as we use here has a chance performance rate of 1/3. If we were interested in estimating some “threshold” of a psychometric function, the standard logistic link function might be considered inappropriate for these data: it could predict that performance falls below 0.33, which if it occurs in observed data can only be due to measurement error or to observers incorrectly switching responses (and is therefore not a desirable prediction to make in general; though see our third caveat in the Discussion). However, we are not estimating thresholds here, but rather we wish to quantify performance differences between discrete levels and also the extent to which performance is different to chance performance. The standard logistic link function is therefore more desirable.

